# Causes and consequences of bacteriophage diversification via genetic exchanges across lifestyles and bacterial taxa

**DOI:** 10.1101/2020.04.14.041137

**Authors:** Jorge A. Moura de Sousa, Eugen Pfeifer, Marie Touchon, Eduardo P.C. Rocha

## Abstract

Bacteriophages (phages) evolve rapidly by acquiring genes from other phages leading to mosaic genomes. Here, we identify numerous genetic transfers between distantly related phages and aim at understanding their frequency, consequences and the conditions favoring them. Gene flow tends to occur between phages that are enriched for recombinases, transposases and non-homologous end joining, suggesting that both homologous and illegitimate recombination contribute to gene flow. Phage family and host phyla are strong barriers to gene exchange, but phage lifestyle is not. We observe more exchanges between temperate phages even if they tend to have smaller genomes. These acquisitions often include transcription regulators and lysins. Yet, there is also extensive gene flow between temperate and virulent phages, or between the latter. These predominantly involve virulent phages with large genomes previously classed as low gene flux, and lead to the preferential transfer of genes encoding functions involved in cell energetics, nucleotide metabolism, DNA packaging and injection, and virion assembly. Such exchanges may explain the acquisition of genes in virulent phages, which tend to have the largest genomes. We used genetic transfers, which occur upon co-infection of a host, to compare phage host range. We found that virulent phages have broader host ranges and mediate genetic exchanges between narrow host range temperate phages infecting distant bacterial hosts, thus contributing to gene flow between virulent phages, as well as between temperate phages. This gene flow drastically expands the gene repertoires available for phage and bacterial evolution, including the transfer of functional innovations across taxa.

## Introduction

Bacterial viruses (bacteriophages or phages) are ubiquitous across environments. Their genomes vary considerably in size, from less than ten genes, up to hundreds (Hatfull and Hendrix 2011; Zhan and Chen 2019; Callanan et al. 2020). Double-stranded DNA phages have by far the largest genomes and are usually regarded as the most abundant (Dion et al. 2020). They can be either virulent or temperate. Virulent phages produce lytic infections, where rapid viral replication ends in progeny release and bacterial death. Temperate phages typically follow either a lytic or a lysogenic cycle (St-Pierre and Endy 2008). The viral DNA remains as a prophage in lysogens, with most genes silent until a signal activates their lytic cycle. Yet, a few progenes may be expressed and provide adaptive phenotypes to their hosts (Brüssow et al. 2004; Harrison and Brockhurst 2017). Lysogeny is highly relevant for the evolution of bacteria, since half of the bacterial genomes have at least one prophage and some have up to 20 prophages (Touchon et al. 2016). Phages also drive horizontal gene transfer among bacteria by transduction (Touchon et al. 2017), which may disseminate virulence factors (Penadés et al. 2015) and antibiotic resistance (Fillol-Salom et al. 2019).

The high genomic plasticity of temperate lambdoid phages has been extensively studied (Botstein 1980; Hendrix et al. 2000; Hatfull and Hendrix 2011), revealing patches of regions of very similar sequences within pairs of very dissimilar genomes. This mosaicism is facilitated by the modular organization of phage genomes and by the role of recombination in the production of phage genome concatemers that are packaged into the virion (Smith 1983). The molecular mechanisms underlying the mosaicism may involve phage-encoded recombinases, which are more permissive to differences between sequences than bacterial RecA-mediated homologous recombination (Martinsohn et al. 2008; De Paepe et al. 2014). Acquisition of genes may also be facilitated by DDE recombinases (*e.g.* insertion sequences (Siguier et al. 2014)), even though they are thought to be rare in phages (Leclercq and Cordaux 2011), or involve homology-free mechanisms such as non-homologous end joining or other types of illegitimate recombination (Morris et al. 2008; Bertrand et al. 2019).

The evolutionary dynamics of phage genomes reflect their distinct lifestyles. A recent study showed that phages can be classed in two “evolutionary modes” representing the relative importance of genetic exchanges in their evolution (Mavrich and Hatfull 2017). High gene content flux (HGCF) phages are 21% of the total and acquire and lose genes at much higher rates than the remaining low gene content flux (LGCF) phages. Most HGCF phages (80%) are temperate and most LGCF phages (87%) are virulent. This extensive mosaicism of temperate phages may be caused by genetic exchanges between phages co-infecting the cells, between prophages in poly-lysogens, or between prophages in infecting temperate phages (Hendrix et al. 2000; De Paepe et al. 2014). The prevalence of prophages in bacterial genomes provides ample opportunity for such exchanges. There is little information on how virulent phages acquire novel genes since they make the vast majority of LGCF phages. Yet, the size of some very large virulent phages (Mesyanzhinov et al. 2002) suggests they have acquired genes from other genomes, and previous studies revealed genetic exchanges in T4-like and in T7-like phages (Filée et al. 2006; Dekel-Bird et al. 2013). Opportunities for genetic exchanges involving virulent phages might be rare, and even less is known regarding genetic exchanges between temperate and virulent phages, even if they are also assumed to be rare. For instance, a recent study found no evidence of recent exchanges between a set of 84 virulent and temperate phages of *Eschericha coli* (De Paepe et al. 2014), and a broader analysis suggested that temperate phages rarely have extensive homology to virulent phages (Mavrich and Hatfull 2017). There are reports of virulent phages acquiring genes from temperate phages or their prophages (Bouchard and Moineau 2000; Garneau et al. 2008), but many concern pairs of closely related phages where one has recently lost the ability to lysogenize its host (Ford et al. 1998; Lucchini et al. 1999; Durmaz and Klaenhammer 2000; Schuch and Fischetti 2006). Genetic exchanges between distantly related phages are likely to contribute to the diversification of their gene repertoires, given the overall dissimilarity of their genomes. Yet, little is known regarding these types of exchanges.

Here, we systematically identify gene transfers between distantly related phages and associate these transfers with host clade, phage family, specific protein function, recombination mechanisms and phages’ lifestyles. Genetic transfers between cellular organisms are usually detected from variations in the density of polymorphism in genomes or the phylogenetic congruence of core genes (Martin et al. 2011). This is impossible when analyzing divergent phages because they have no core genes that could be used to build reliable phylogenies and many genes lack homologs in other phages. Furthermore, the extensive mosaicism of certain temperate phages implies that most genes have different phylogenies. Therefore, we searched instead for strong mosaicism, i.e. for a few highly similar genes within highly dissimilar genomes. We were able to identify many events of gene transfer, even if our conservative approach will miss very ancient transfer events, or those that occur between similar genomes. Our findings show that functionally diverse genes are transferred between distantly related phages, both within and between phage lifestyles, regardless of their described gene flux mode. Moreover, our results suggest that virulent phages facilitate the transfer of genes across more distant taxa of temperate phages. This increases the repertoires of genes available for both phage and bacterial evolution.

## Results

### The network of similarities between phages

We analysed a dataset of 2487 complete phage genome sequences and used PHACTS (McNair et al. 2012) to predict their lifestyle, having identified 1161 virulent and 1336 temperate phages. To control for the difficulties in phage lifestyle assessment, we repeated all the key analyses of this study while restricting the phage dataset to the high confidence predictions of PHACTS, and using the alternative tool BACPHLIP (Hockenberry and Wilke 2020), which uses an extensive manual curation of the data from (Mavrich and Hatfull 2017)). We systematically found similar qualitative results for these controls. They are only briefly mentioned in the main text, but are presented in detail in the supplementary material. Table S1 describes the controls for the main findings and guides through the associated supplementary material. For this study, we excluded the phage genomes with less than 15 predicted proteins (110 genomes), as mosaicism cannot be reliably identified using our method (see below). Many of the excluded phages correspond to ssDNA or ssRNA phages. We then searched for reciprocal best hits between pairs of phages and computed the weighted Genome Repertoire Relatedness (wGRR, see Materials and Methods) for the remaining 2387 phage genomes (1297 temperate and 1090 virulent, 99.9% of the total being dsDNA phages). This resulted in 2,847,691 pairwise wGRR values, among which ca. 91% were null (no detectable homology). Still, 99.9% of the phages had homologs in at least one phage. The histograms of non-null wGRR values (Fig 1A, see also Fig S1A and C for the distributions obtained using alternative lifestyle classifications) revealed higher values for the comparisons between pairs of phages with similar lifestyles (temperate-temperate or virulent-virulent) than for the comparisons involving virulent and temperate phages. Nevertheless, we found clear homologs in ~6% of the comparisons between phages with different lifestyles, even if the wGRR values were small in most cases. This reveals a network of homology across almost all dsDNA phages, even across lifestyles.

**Fig 1.**
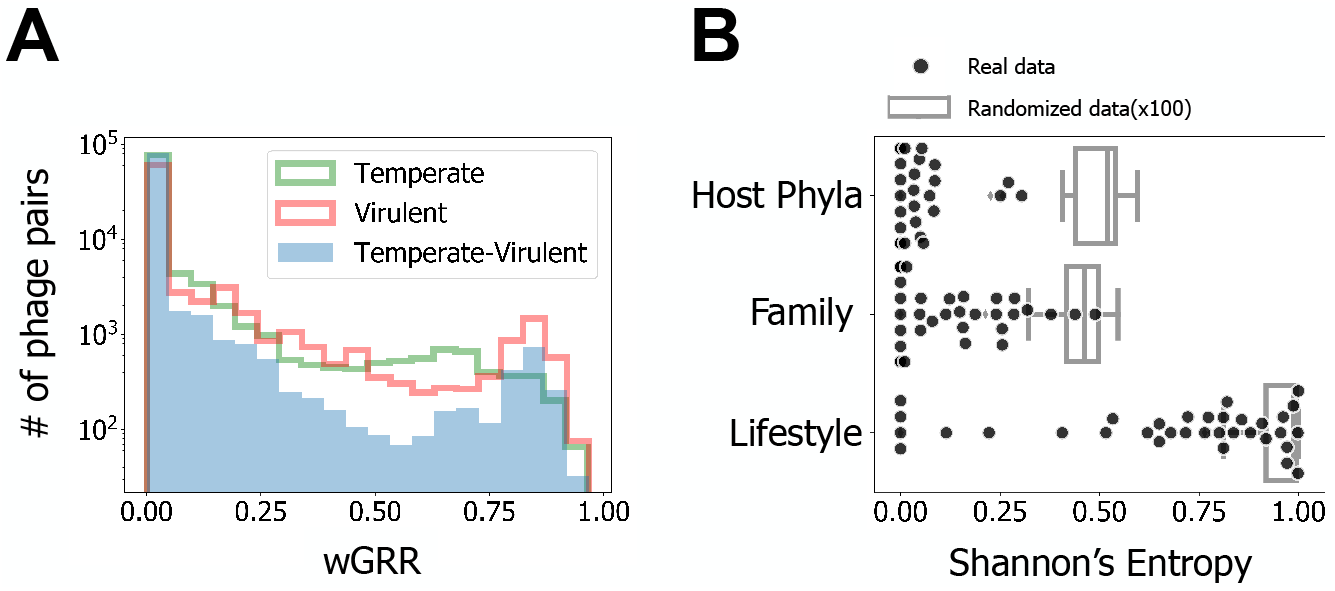
Results of the analysis of similarity between phages. A) Histograms of the wGRR values (with wGRR>0). B) Shannon’s Entropy values for each cluster identified with the Louvain community detection method in the wGRR matrix. Results are given for the three phage traits (N=34 for each trait, one per cluster). Boxplots represent the distribution of the concatenation of 100 repetitions of a process where phage lifestyles, hosts and families are randomly assigned to each node (N=3400 for each trait). All distributions show significantly lower values (less heterogeneous clusters) than their random counterparts (all p<0.0001, 2-sample Kolmogorov Smirnov test). The observed clusters have significantly higher values (more heterogeneous clusters) when analyzed in terms of phage lifestyle than in terms other variables (both p<0.001, Tukey HSD Test).

To analyse the relations of homology between phages, we built a graph where phages are nodes and edges represent wGRR values (Fig S2). These networks are frequently used to describe evolutionary relationships between phages (Lima-Mendez et al. 2008; Halary et al. 2010). We then clustered the phages by their wGRR using the Louvain method for community detection. This resulted in 34 clusters with at least 3 phages, 5 with 2 phages, and 3 singletons (Fig S3). In order to compare the effects of the phage lifestyle (virulent or temperate), family (the characterized virion morphology) and host phyla in the separation of phages in different clusters of wGRR, we used Shannon information entropy. This index quantified the homogeneity of each cluster regarding each trait (Fig 1B, S2B and D). Clusters are very homogeneous in terms of host phyla and phage family, as revealed by average entropies close to zero (average Shannon Index of 0.05 and 0.13 respectively). In contrast, they are significantly more heterogeneous regarding their lifestyle (average Shannon index of 0.67, p<0.001, Tukey Honest Significant Difference (HSD) test), and almost all clusters included both temperate and virulent phages. These results show that phages from different families and bacterial phyla tend to be well separated, whereas there is significant genetic similarity between many temperate and virulent phages independently of phage or bacterial taxonomic considerations.

### Recent genetic transfer across distantly related phages

Sequence similarity between phages can be explained by common ancestry or by recent gene flux. When the wGRR is small, the two processes translate into different patterns: common ancestry results in many homologs of low similarity along the genomes, whereas recent genetic transfers result in a small number of highly similar homologs within very dissimilar genomes. Note that we consider that a genetic transfer corresponds to the transfer of one or several genes from one phage to another. To distinguish between recent transfers versus ancestry one can use the contrast between the fraction of homologous genes (above a low minimal similarity threshold of 35% identity) and the fraction of genes that are very similar (more than 80% identity). Distant common ancestry results in a sizeable fraction of the pairs of genomes with homologs but no single very similar gene, whereas recent gene transfers between distantly related phages result in a few highly similar homologs and a low wGRR. This procedure does not allow to identify genetic transfers that are very ancient, that occur between very closely related genomes or that involve a large fraction of the genome. However, since we showed above that most comparisons between phage genomes have low wGRR values, these cases should be rare when comparing distantly related phages (Fig 1A). Indeed, comparisons between temperate and virulent (Fig 2A and S4C) and between virulent phages (Fig 2B and S4B) show a few nearly identical genes (>80% identity) within genomes otherwise lacking homologues. These cases are best explained by recent genetic transfers. In contrast, when genomes exchange genes very frequently, as is the case of many temperate phages, the comparisons show mosaics of highly similar and highly dissimilar (eventually non-homologous) genes. This results in a linear distribution where almost identical genes co-exist in very different genomes (Fig 2C and S4A).

**Fig 2.**
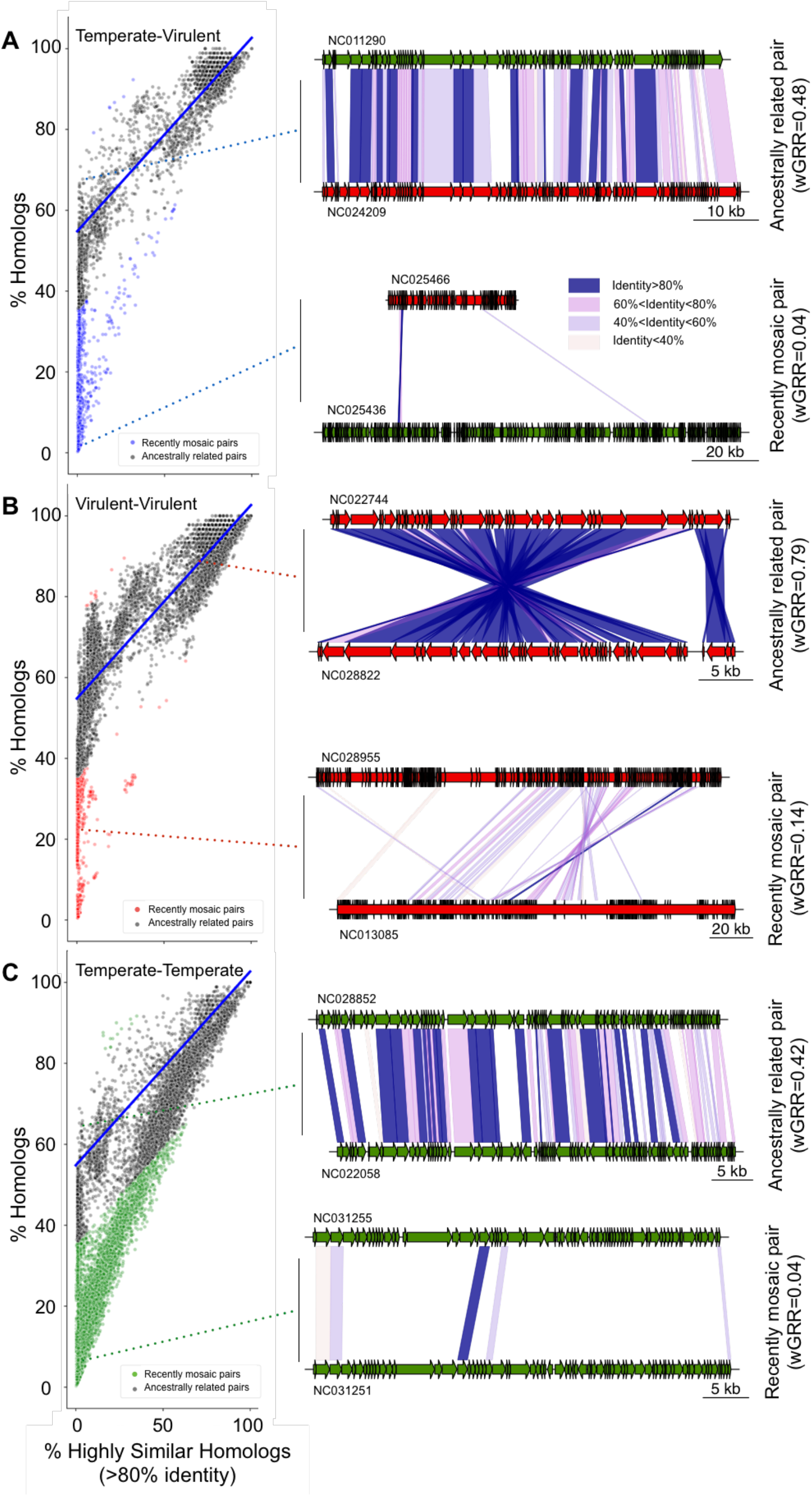
Identification of putative phage pairs with recent gene transfers. A-C) Scatterplots of pairs of phage genomes, with recently mosaic pairs indicated as colored points (otherwise grey). The linear regression model (blue lines) was inferred for the temperate-virulent dataset in A and applied to all datasets. The genomic maps show representative examples of one recently mosaic (bottom) or one ancestrally related phage pair (top), for each phage lifestyle combination (see more examples in Fig S7). Dashed lines indicate the location of the phage pairs in the distributions on the left. Color codes for points and genes: virulent (red), temperate (green), virulent-temperate (blue). Colors in the blocks linking the phages indicate sequence similarity between homologs.

We then built a linear model based on the relationship between the fraction of very similar proteins (>80% identity) and the fraction of homologous proteins between temperate and virulent genomes (Fig 2A, blue line). We limited this dataset to pairs of phages with at least 50% of homologous proteins, where the influence of outliers that are associated with recent genetic transfers is expected to be weaker. The linear model fitted to the temperate-virulent dataset represents a conservative model of the null hypothesis that the relationship between these two parameters is mostly due to ancestry. It fits well the major group of comparisons between virulent and temperate phages across almost all the range of the regression (Fig 2A). We computed the negative residuals of the linear model to identify significant negative deviations to the main trend. These represent cases where genomes have an unexpectedly high number of very similar genes given the overall level of homology (Fig 2). This threshold on the value of negative residuals was used across all datasets to identify putative recent events of genetic transfer. For pairs of virulent-virulent or temperate-virulent phages, the overwhelming majority of exchanges were inferred between very dissimilar phages (wGRR below 0.2, Fig 3 and S5). The comparisons between temperate-temperate phage pairs have a notably different distribution of genetic similarity (Fig 3B, S5B and D), with a large fraction showing intermediate (>0.25) wGRR values. This is suggestive of higher gene flux in temperate phages, in agreement with previous works (Hatfull and Hendrix 2011; Dion et al. 2020). For simplicity, phages inferred to be involved in recent genetic transfers with other distantly related phages will from now on be referred as “recently mosaic phages” or “phages with recent genetic transfers” (even if we do not presume that genetic transfers that lead to mosaicism do not occur between closely related phages).

**Fig 3.**
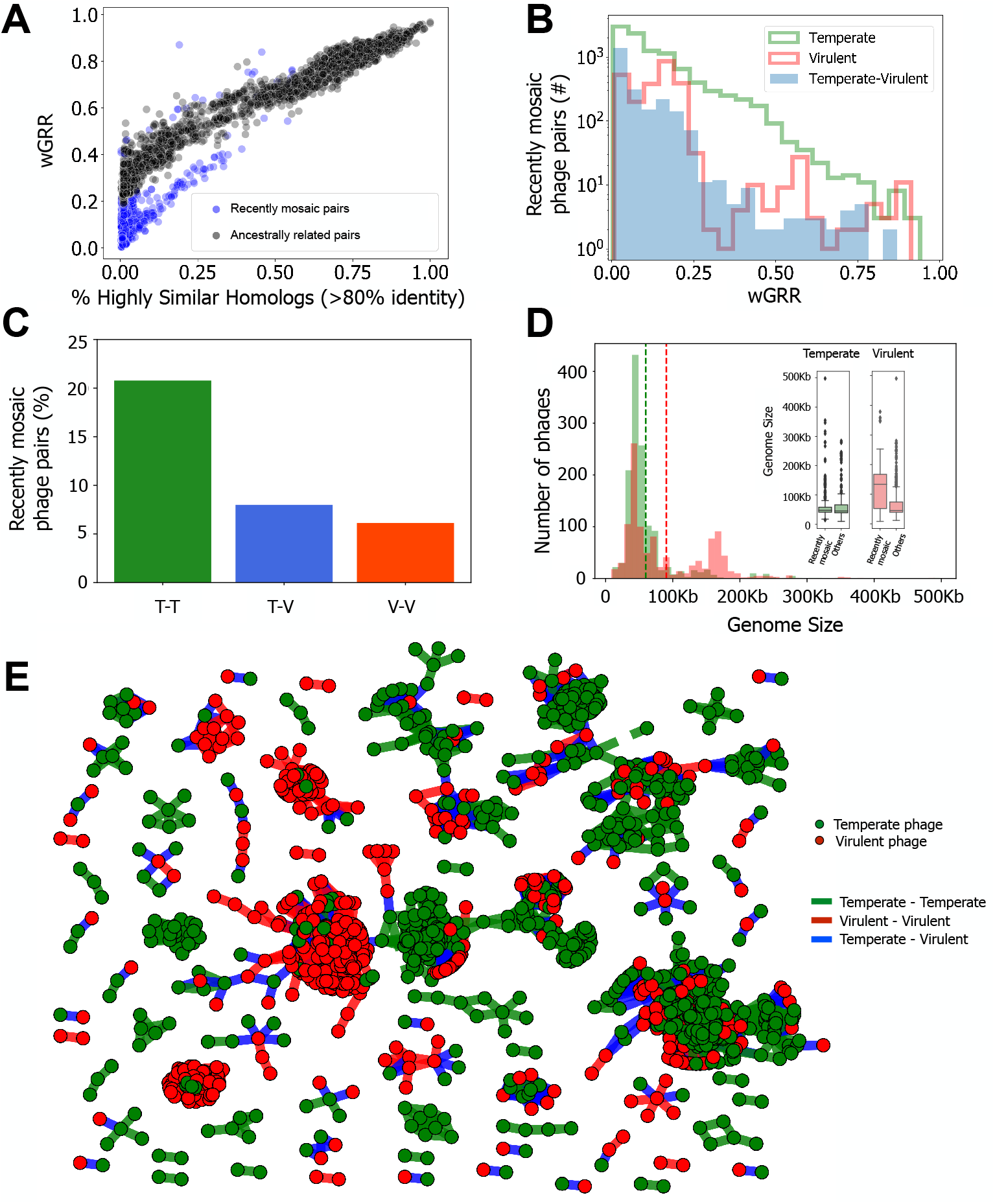
Identification of phage pairs with putative gene transfers in the wGRR network. A) Scatterplot of pairs of temperate-virulent phages in terms of wGRR and the fraction of high sequence-identity homologous genes. B) Histogram of wGRR values for the subset of recently mosaic phage pairs. C) Frequency of recent genetic transfers within or between lifestyles, for pairs of phages with wGRR>0.01 (p<0.001 for association of genetic transfers with the phages’ lifestyle, χ^2^ test). T-T: temperate-temperate, T-V: temperate-virulent, V-V: virulent-virulent. D) Distribution of the sizes of genomes of temperate (green) and virulent phages (red). Dashed lines indicate the average genome size for each lifestyle (in green, temperate phages, 59Kb; in red, virulent phages, 90Kb). In the inset, the boxplots show the distribution of genome sizes for the phages involved in recent transfers versus the genomes of the other phages, for each lifestyle. E) Network of recently mosaic phages (i.e., those with wGRR<0.5 in panel B). Each node represents a phage genome and each edge a relationship of genetic similarity. Temperate phages are shown in green nodes, virulent phages are shown in red nodes, and the edge colors correspond to events of gene transfer between temperate-temperate (green), temperate-virulent (blue) and virulent-virulent (red) phages.

### Gene flux in relation to genetic and lifestyle differences

The analysis of sequence similarity between proteins of phages connected in the wGRR network shows two distinct patterns: genomes that are more likely to be ancestrally related tend to have large regions of low sequence similarity, whereas the others show homology limited to one or a few very similar genes (Fig 2 and S7). This suggests that our method discriminates recent genetic transfers from shared ancestry. 48% of the virulent phages (528 out of 1090) and 66% of the temperate phages (852 out of 1261) were among the recently mosaic phages, being involved in at least one, but potentially more, events of recent genetic transfers.

Phage genomes can vary by orders of magnitude in size, and one would expect larger genomes to have more accessory genes and thus engage more often in gene acquisition. Yet, even though genomes of temperate phages tend to be classified as HGCF, their genome size is significantly smaller than the one of virulent phages (59 vs 90kb, p<0.001 2-sample Kolmogorov-Smirnov, Fig 3D). Within temperate phages, the recently mosaic are slightly smaller than the others (inset of Fig 3D). In contrast, recently mosaic virulent phages have much larger genomes than the others (115 vs 67kb, p<0.001 2-sample Kolmogorov-Smirnov, inset of Fig 3D). The analysis of genetic transfers between unrelated phages of different lifestyles showed that 36% of the virulent phages were inferred to transfer genes with temperate phages, whilst 37% of temperate phages were found to transfer genes with their virulent counterparts (at least once). This means that transfers are divers, *i.e.*, there is not a small cluster of one type of phages responsible for all the transfer with a variety of phages of another type. The network of wGRR values restricted to recently mosaic phage pairs (Fig 3E) confirms that clusters are very homogeneous in terms of host phyla or phage family, but often put together phages with different lifestyles (Fig S8A-B, p<0.001, Tukey HSD test, see also Figs S8C-D and S9). This further substantiates the previous results showing that among the three different ways of partitioning the phages, two of them define strong barriers to genetic transfers – families and bacterial clades – whereas lifestyle does not. Importantly, it shows that the co-occurrence of virulent and temperate phages in clusters of the network is partly driven by their genetic exchanges.

As expected, genetic transfers depend on the genetic relatedness and on the lifestyle (Fig 3B-E). They are also much more frequent between temperate phages, being observed in ~21% of the temperate-temperate pairs (from all that have wGRR>0.01). The frequency of transfers between pairs of virulent and temperate phages is smaller (~8%) and almost as low as that among virulent phages (~6%, Fig 3C, Fig S10). To analyse the relative roles of wGRR and lifestyle in shaping the frequency of genetic transfers, we excluded from the dataset the pairs of phages with very low wGRR values since these are very numerous and almost never reveal recent transfers (in two separate analyses, either wGRR>0.01, Fig 3C, or wGRR>0.05, Fig S10). We made a stepwise logistic regression using the remaining data where the dependent (binary) variable was the presence of genetic transfers and the independent variables were the wGRR and the lifestyle of the pair. Both variables contributed significantly to explain the variation in the frequency of genetic transfers (Table S2). The contrast between pairs of temperate phages and the remaining pairs had the highest contribution to the logistic regression, followed by the wGRR. This analysis shows that the frequency of recent genetic transfers is dependent on the lifestyle, even when accounting for genetic distances between phages.

We then investigated whether genomes classed as HGCF drive the genetic exchanges between dissimilar phages. We matched our database with that of (Mavrich and Hatfull 2017) and classified 283 phages as HGCF and 737 as LGCF (ca. 43% of our dataset). Pairs of HGCF phages had more cases of recent genetic transfers (8%) than pairs of LGCF phages (4%) (p<0.001 Fisher exact test). This trend was driven by temperate phages, where HGCF pairs of recently mosaic phages are much more frequent than LGCF ones (2998 vs 89). However, the vast majority (>95%) of the recent genetic transfers involving virulent phages (either with other virulent phages or with temperate ones) occurred between LGCF phages (Fig S6). This is possibly because few virulent phages were classed as HGCF. The absolute number of recently mosaic pairs of LGCF virulent phages (1119) or LGCF virulent-temperate phages (478) is higher than the number of HGCF pairs in both cases (3 and 87 respectively). Hence, HGCF phages drive a disproportionally high number of genetic transfers in temperate phages, but they do not account for the majority of the transfers that involve virulent phages.

### Functional classification of recent genetic transfers

The identification of pairs of recently mosaic phages allows to study the actual genes involved in these transfers. We defined them as those encoding highly similar proteins (at least 80% identity) in two dissimilar phage genomes (wGRR lower than 0.5). We found an average of 6 recently transferred genes per phage genome. These genes were only slightly smaller on average than the other genes (191 vs 207 amino acids, Fig S11). Since gene flow between genetically dissimilar phages can contribute to their functional diversification, we analysed the functions of these recently transferred genes in light of phage and bacterial biology. We clustered the Prokaryotic Virus Orthologous Groups (pVOGs) profiles database into functional classes and used it to associate genes involved in genetic transfers with phage-related functions (Fig 4A, Fig S12, Table S3 and File S2). The set of recently transferred genes matched a very large number of different and diverse protein profiles (>100) even if most are of unknown function. This shows that the genes transferred between distantly related phages are not restricted to a few functional categories of slow evolving proteins that might spuriously be identified as transfers. Some functions are overrepresented in transfers that occur between virulent-virulent or temperate-virulent pairs of phages (e.g., proteins involved in packaging, injection and assembly), but underrepresented in temperate-temperate exchanges. On the other hand, proteins associated with lysis are more often transferred between temperate phages and rarely between virulent ones.

**Fig 4.**
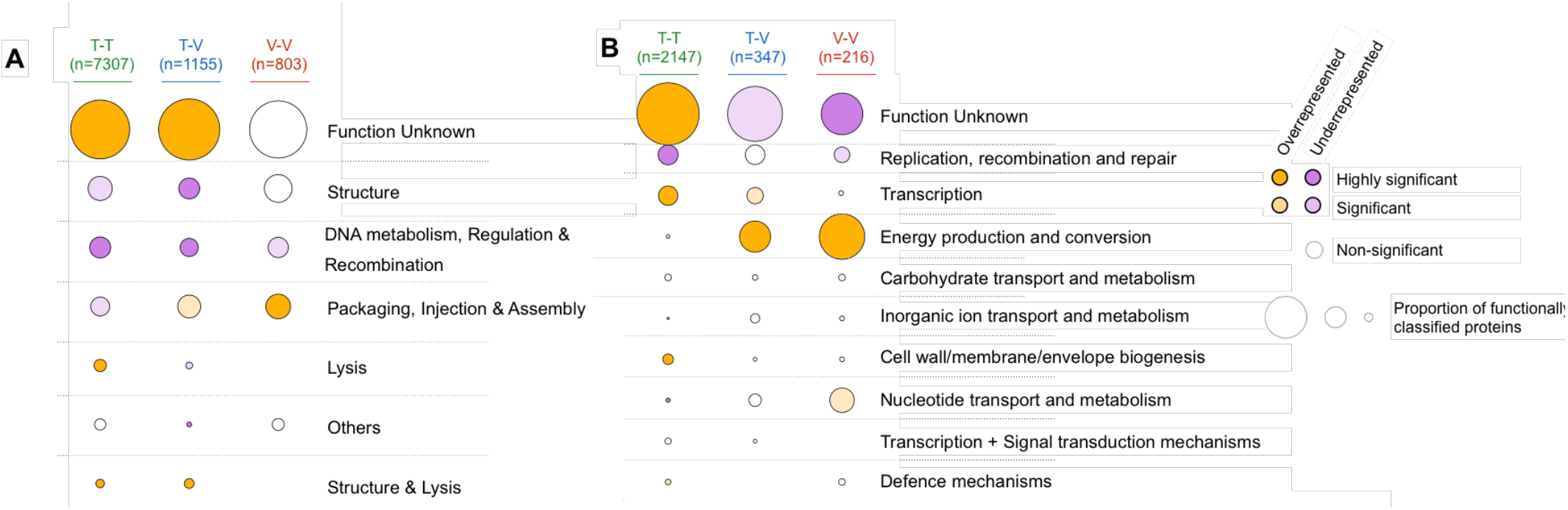
Functional classification of genes involved in genetic transfers between distantly related phages. **A)** Classification of typical phage functions using the pVOG database. **B)** Classification of typical bacterial functions using the bactNOG database. In both panels, the size of each circle corresponds to the proportion of genes with a given function. The total number of genes with an assigned function is shown at the top. Enrichment (orange) or depletion (purple) of each function is assessed relatively to classification of all phage proteins with pVOG (in **A**) or bactNOG (in **B**). In both cases, the color intensity corresponds to the statistical significance of the classification (Fisher-exact test adjusted for multiple comparisons with the Benjamini-Hochberg method).

Temperate phages sometimes carry genes that were acquired from bacterial genomes, which also leads to the possibility of these genes being transferred between phages. We thus searched if genes identified as genetic transfers were associated with some specific bacterial functions. We removed from the bacteria Non-supervised Orthologous Groups (bactNOG) profiles database the functions associated with phages by removing profiles matching pVOG profiles with a viral quotient (VQ) higher than 80%. We then used the remaining bactNOG profiles to annotate the transferred genes (Fig 4B, Fig S12, Table S4 and File S2). The vast majority of these genes have no defined function. Some of the remaining might be typical phages genes that were missed by the filter using the pVOG profiles. Yet, other genes matched profiles whose functions are related to carbohydrate or nucleotide transport and metabolism, DNA related functions, or energy production and conservation. The latter category is particularly overrepresented in virulent-virulent and temperate-virulent transfers, and in both cases, this category represents the majority of genes with a known assigned bactNOG function, consisting of proteins implicated in photosynthetic activity in phages infecting marine bacteria. In contrast, genes encoding these functions were rarely detected as being recently transferred between temperate phages. Other notable bacterial functions transferred between distantly related phages include methylases, transporters and single-strand binding proteins. Hence, our results suggest that genetic transfers between unrelated phages facilitate their functional diversification and the dissemination of bacterial traits.

### Phages with recent genetic transfers are enriched for mechanisms of genomic exchanges

Genetic transfers between distantly related phages cannot usually be achieved by the cellular RecA recombinases, because these tolerate few mismatches (Shen and Huang 1986). However, some phage recombinases, e.g. lambda Red from the RecT family, allow recombination between more dissimilar sequences (De Paepe et al. 2014). To test if these recombinases could facilitate genetic transfers between distant phages, we searched for the Erf, Sak, Sak4, RecT, UvsX and Gp2.5 families in the genomes of phages. We found that ca. 42% of the phages with recently transferred genes encode at least one of these recombinases, while only 26% of the remaining phages encode them (Table S5). This suggests that the presence of these recombinases may favour genetic transfers between unrelated phages, as previously shown in several *E. coli* phages (Martinsohn et al. 2008; Hutinet et al. 2018).

The distribution of the different types of recombinases differs between phage lifestyles (Fig 5, Fig S13 and File S3). Recently mosaic temperate phages are enriched for all recombinases tested, relative to the other temperate phages, with a particularly striking over-representation of RecT-like recombinases. Recently mosaic virulent phages are particularly enriched for one specific recombinase – UvsX – that is present in 70% of them. We then set to explore the possibility that phages classed as HGCF have more recombinases. The latter are present in most (62%) temperate HGCF phages, but only in a few LGCF temperate phages (27%). The same analysis showed no remarkable difference between virulent phages (Table S6). This suggests that phage recombinases can underlie the higher gene flux previously detected in HGCF phages.

**Fig 5.**
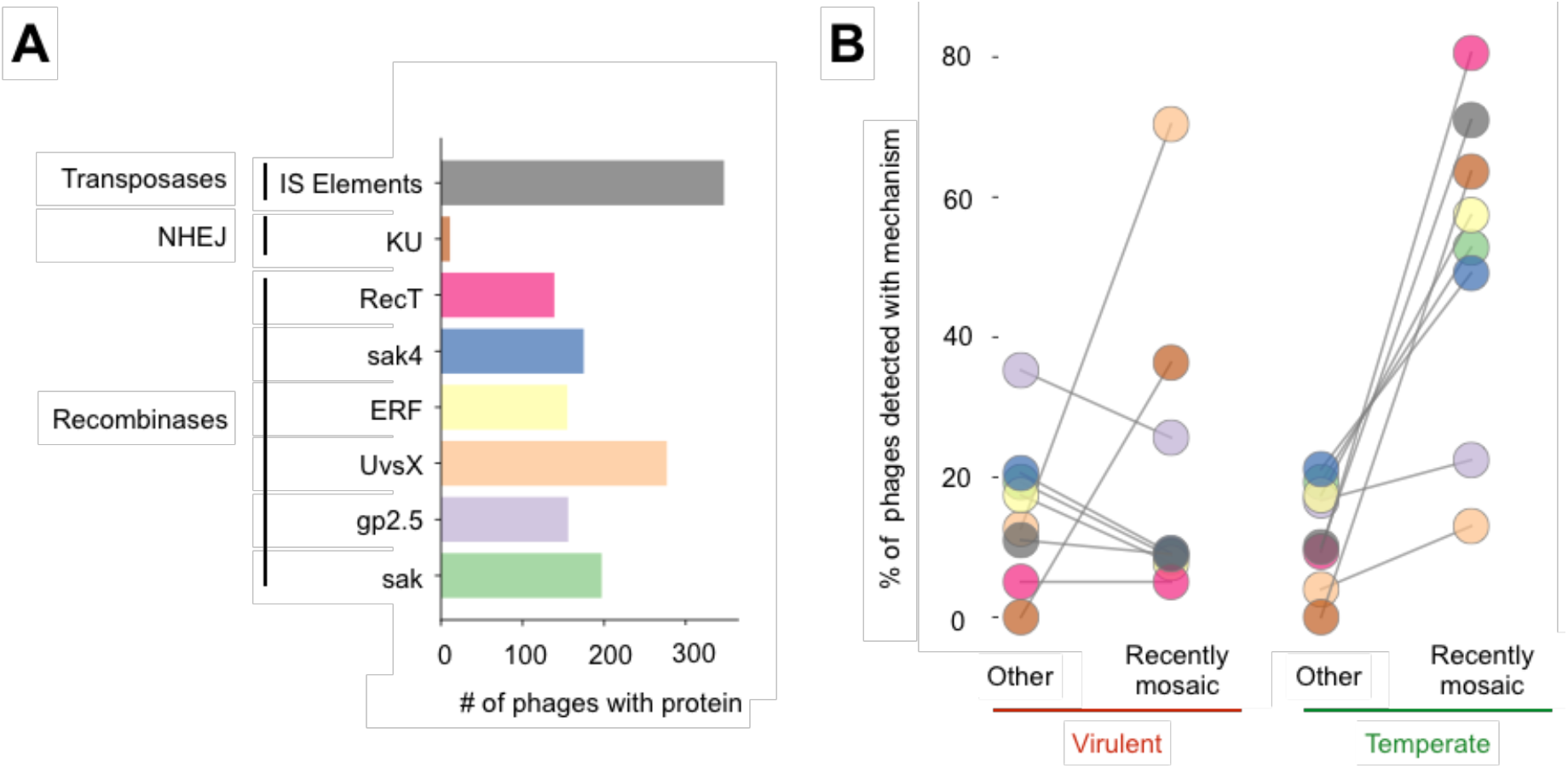
Putative mechanisms involved in genetic transfers and their relative frequency in recently mosaic phages. **A)** The total number of phages with at least one homologous gene for each of the proteins’ types. **B)** Proportion of genomes with homologous genes for each type of proteins analysed in recently mosaic phages and in the remaining phages. Colors of the bars and circles correspond to the different types of proteins analysed.

Genetic transfers between phages can only occur within bacterial hosts. Phage-like recombinases encoded in the bacterial genomes may thus play a role in these transfers. We searched for these recombinases in the bacterial hosts, discarding UvsX recombinases from the analysis, since they are part of the RecA family that is encoded by most bacterial genomes (Rocha et al. 2005). We found that the hosts of phages with recent genetic transfers are 29% more likely to encode recombinases in their genomes than the hosts of the other phages (0.75 vs 0.58 recombinases on average, across all genomes available for each host). These host recombinases are concentrated (ca. 90% of the total) in prophages and absent from the rest of the bacterial genomes (Table S7). This suggests that recombinases from prophages facilitate gene transfers with other phages within bacterial cells. In agreement with this view, the genomes of bacterial species that host phages with recent transfers tend to have ca. 29% more prophages (2.2 and 3.1 prophages on average, for the hosts of temperate and virulent phages respectively) than the remaining phages (p<0.001, two sample Kolmogorov-Smirnov test, Fig S14-S15).

Genetic transfers between phages may also result from non-homologous or illegitimate recombination (Morris et al. 2008). Many mechanisms of illegitimate recombination lack specific genes that can provide an indication of its frequency in a genome. However, the non-homologous end-joining (NHEJ) pathway is a very well characterized mechanism that can resect DNA double strand breaks lacking homology by the action of a very specific protein called Ku (Aravind and Koonin 2001). We searched for genes encoding this protein and found that bacterial genomes that are hosts of phages with recent genetic transfers are 13% to 26% more likely to encode NHEJ than the others (p=0.002 for temperate and p<0.001 for virulent phages, two sample Kolmogorov-Smirnov, Fig S16). Recently, a Ku homolog in the Mu phage, was shown to promote NHEJ using a host ligase (Bhattacharyya et al. 2018). Interestingly, we found 11 phages in our dataset (7 temperate and 4 virulent) encoding homologs of the Ku protein (Fig 5, Fig S13 and File S3). All of these phages were involved in recent genetic transfers. Even if their number is small, the over-representation of phages carrying Ku within those with recently transferred genes is significant (p=0.003, Fisher exact test).

Finally, transposase-mediated gene transfers can also occur between unrelated genomes. We found 347 temperate phages coding for insertion sequences (IS). Around 71% of these are recently mosaic temperate phages, which is more than expected by chance among these phages (p<0.001, Fisher exact test, Fig 5, Fig S13 and File S3). We also found that 19 temperate phages have more than 3 IS in their genomes (up to 18, in one phage genome). In contrast, transposases are very rare in virulent phages, suggesting that temperate phages tend to acquire them when they are integrated in the bacterial chromosome. There is no difference in the number of transposases between virulent phages with recent genetic transfers and the others (p=0.71, Fisher exact test). Overall, our results indicate that recently mosaic phages, as well as their hosts, are enriched for different mechanisms that potentiate the transfer of genetic material across distantly related phage genomes.

### Broader host range of virulent phages facilitates genetic transfers across clades of temperate phages

Defining the host range of phages is a notoriously difficult problem. However, if recent genetic transfers are observed between phages that have distinct bacterial hosts, it is reasonable to assume that either their host ranges overlap or that their recent ancestors must have at some point infected the same bacteria. Consequently, tracking these genetic transfers provides information about the phages’ host range. We computed both the patristic distance (based on the 16S rDNA, Fig 6A, S17A and C) and the differences in 3-mer genomic signatures between the bacterial hosts (identified at the species level) of all pairs of phages with recent genetic transfers (Fig 6B and S17B and D, see also Fig S18). In all three datasets (transfers between temperate-temperate, between virulent-virulent and between temperate-virulent), most cases of transfers occurred within very closely related bacteria. However, the datasets significantly differ in their host range (p=0.001, Tukey HSD test). Importantly, pairs of virulent phages with recently transferred genes are more frequently associated with distant (or dissimilar) hosts, with an average patristic distance of 0.08, than pairs of temperate phages (average patristic distance of 0.03, p=0.001, Tukey HSD test). We analysed in more detail the few cases of transfers between pairs of temperate phages with very distantly related bacterial hosts (outliers in Fig 6, see also File S4). They involve either temperate phages isolated in high temperature environments, or concern phages classed as temperate by PHACTS but virulent by BACPHLIP. Overall, our results suggest that virulent phages are able to infect (and transfer genes within) a wider range of host species than temperate phages. Interestingly, genetic transfers between temperate and virulent phages are associated with intermediate distances between bacterial hosts (average patristic distance of 0.05), which likely reflects the contribution of broader host (virulent) and narrower host (temperate) phages.

**Fig 6.**
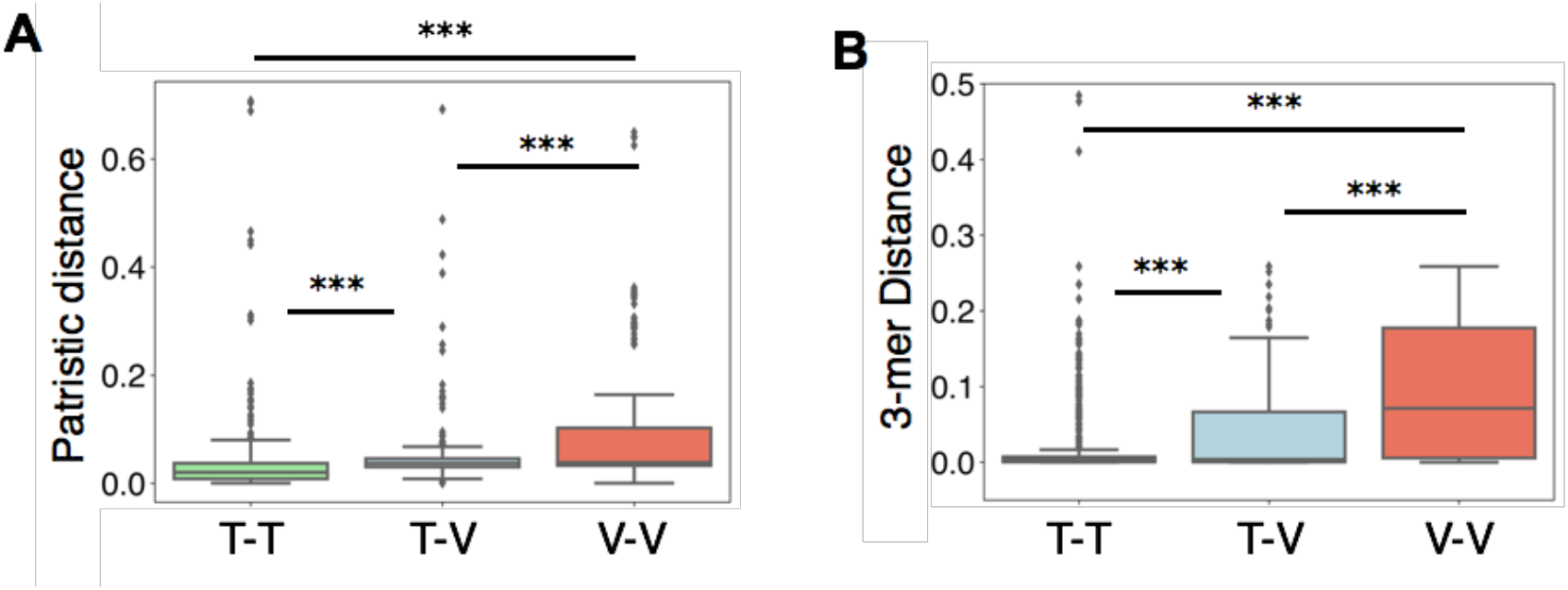
Recent genetic transfers between phages indicates a broader host range of virulent phages. **A)** Boxplots of the non-null patristic distances (computed from the 16S rDNA gene tree) between the hosts’ species of each recently mosaic phage pair. **B)** Distributions of the differences in 3-mer genomic signatures between the hosts’s species of each recently mosaic phage pair. *** p=0.001, Tukey HSD for all pairs. T-T: temperate-temperate, T-V: temperate-virulent, V-V: virulent-virulent.

If virulent phages have broader host ranges and transfer genes from/to temperate phages in phylogenetically distant bacteria, then they can potentially shuttle genes between those temperate phages that are sexually isolated (because they infect distantly related bacterial clades). To systematize these observations, we built a simplified network only representing nodes and edges between temperate-virulent phage pairs with recent genetic transfers, and only when there are two temperate phages (each with a distinct host’s genus) that are “linked” by a virulent phage (Fig 7A and S19A). This network reveals several transfer events between pairs of virulent and temperate phages from distinct genera, but also indicates that virulent phages can facilitate genetic transfers between temperate phages whose hosts are from distinct genera. In some cases, these transfers concern distinct genes (Fig 7B, S19B-C), whereas in others they concern the exact same gene (Fig 7C, Fig S19D-E). In these examples, temperate phages share little or no homology beyond genes homologous to the virulent phage (Fig S20), suggesting that these transfers were mediated by the virulent phage. Importantly, these results suggest that genetic transfers between temperate and virulent phages, in tandem with the broader host range of virulent phages, increases the frequency of genomic exchanges between temperate phages in phylogenetically distant hosts.

**Fig 7.**
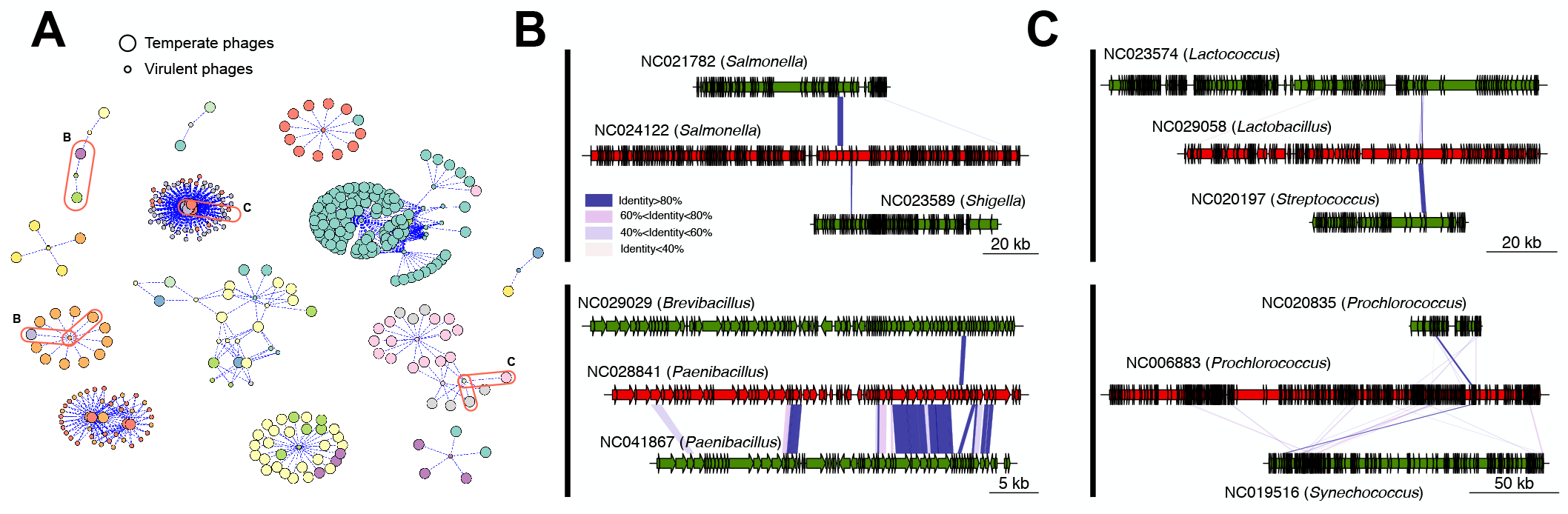
Genetic transfers between temperate phages with different host genera through virulent phages. **A)** Simplified network restricted to pairs of temperate-virulent phages where the virulent phage transferred genes from/to at least two temperate phages infecting distinct bacterial genera (i.e., edges shown are only those that link temperate phages with different hosts’ genera through a virulent phage). Each node color represents a different bacterial host genus, and the node sizes identify either temperate (large nodes) or virulent (small nodes) phages. Examples from panels B and C are highlighted in ellipses. B) Examples of virulent phages (red) with genes transferred from/to two temperate phages (green) infecting distinct bacterial genera. C) Analogous to B, but representing transfers with the virulent phage that involve the exact same protein in both temperate phages. Colors in the blocks linking the phages indicate the level of sequence similarity between homologs. The genomic maps between each of the two temperate phages for each of the cases are shown in Fig S20.

## Discussion

Our study provides the first systematic quantification of how genetic transfers across distantly related phages are shaped by genetic distance, mechanisms of genetic exchange and differences in lifestyle. Our approach has some caveats. First, incorrect classification of lifestyle leads to spurious inference of inter-lifestyle gene transfer events. We controlled for this by using three different procedures of lifestyle classification that revealed similar qualitative patterns, showing that our results are robust to mis-assignment of lifestyles (see Table S1). Second, our method is incapable of identifying certain types of genetic transfers: the ancient ones, those covering a large fraction of the phage genome, and those between closely related phages. It is important to note that this excludes the identification of transfers between temperate phages and their recent virulent variants (e.g., recent loss of an integrase) and it implies that we may have underestimated the number of gene transfers between phages with different lifestyles. Third, the distinction between genetic transfers and ancestry is particularly challenging in the analysis of temperate phages because of their pervasive mosaicism. This leads to less confident identification of specific events of recent gene transfers between temperate phages.

Regardless of these caveats, we could identify in most phages clear traces of recent genetic transfers with distant phages. Transfers between temperate phages can occur between prophages or prophages and infecting phages. But when do the other exchanges occur? We envisage three major scenarios for genetic transfers between virulent or between virulent and temperate phages. First, two virulent phages may infect a bacterium at the same time, or meet in a cell if there is a resident virulent phage in pseudo-lysogeny (Cenens et al. 2013). Second, transfer between phages of different lifestyles may occur when virulent and temperate phages co-infect a bacterium, or when virulent phages infect a lysogen. These transfers can occur in both directions. Virulent phages can acquire DNA from prophages when they degrade the bacterial chromosome, producing many recombinogenic linear double stranded DNA molecules. Temperate phages, or prophages, can acquire DNA from virulent phages when the latter are cleaved by the hosts’ defence systems, such as restriction modification systems or CRISPR-Cas systems, which also results in linear double stranded DNA (Arber 2000; Marraffini 2015). Given the prevalence of bacteria with active or defective prophages, such events may be more likely than those of co-infection. Our approach does not allow to identify the direction of the gene transfers (from temperate to virulent or vice versa), and this is key to know the relevance of the above scenarios. Nevertheless, and independently of the circumstances that facilitate genetic transfers between distantly related phages, this genetic flux is expected to increase when the infected hosts have many prophages.

Which molecular mechanisms mediate gene flux between distantly related phages? Two schools of thought have proposed that genetic transfers between very divergent temperate lambdoid phages could result from either relaxed homologous recombination or illegitimate recombination (Hatfull and Hendrix 2011; De Paepe et al. 2014). These mechanisms are not mutually exclusive, and both were proposed to act on the observed recombination between virulent phages and prophages in *Lactococcus* (Labrie and Moineau 2007). Here, we find evidence for the contribution of both mechanisms in gene transfer across phages, since recently mosaic phages have more recombinases, NHEJ and transposases. However, the relative contribution of these mechanisms depends on the phage’s lifestyles. In virulent phages, gene flow seems associated mostly with UvsX, but in temperate phages it is associated with all the analysed mechanisms. Interestingly, recombinases are among the functions transferred between unrelated phages, suggesting that the potential for further genetic transfers can itself be driven by past exchanges. Hence, genetic exchanges, eventually mediated by recombinases encoded in *cis* or by other mechanisms, facilitate the acquisition of recombinases that will further increase phage genomic plasticity.

In general, our analysis shows a broad diversity of functions exchanged through recent gene transfers, which may facilitate phage adaptation to novel challenges. Indeed, we found that proteins involved in the production of virions, but also bacteriocins or RNA polymerases can be transferred between distant phages. A recent work has also found evidence for the exchange of genes encoding anti-CRISPR proteins between temperate and virulent phages (Hynes et al. 2018), further highlighting the potential for genetic exchanges between unrelated phages to disseminate adaptive traits. Despite the striking functional diversity of the genes involved in such transfers, the functional categories preferentially exchanged depend on the lifestyle of the phages. For example, transfers between temperate phages over-represent genetic regulators, which fits the expectation that these elements require more complex regulation than virulent phages. This difference in functions between lifestyles could also be related with the broader host range we inferred for virulent phages, as functions preferentially transferred into virulent phages might be more effective independently of the host background. The permanence of genes with bacterial functions might be, in most cases, transient in virulent genomes, since they provide no benefit and virulent phages cannot lysogenize their hosts. However, there are examples of auxiliary metabolic genes (Thompson et al. 2011) that have been acquired and repurposed by those phages. Accordingly, the bacterial function we find to be most enriched in the transfers involving virulent phages (either with other virulent phages or with temperate ones) are photosynthetic genes. These genes were shown to increase viral progeny of virulent phages (Mann et al. 2003; Lindell et al. 2004), and were also previously shown to be exchanged between *Prochlorococcus* and *Synechococcus* through viral intermediates (Zeidner et al. 2005). This suggests that gene flow with and between phages plays an active role in the evolution of cyanobacterial photosynthesis. These and other adaptive bacterial traits could further disseminate through transfers that occur between virulent phages, or phages with distinct lifestyles.

Our findings suggest that the impact of genetic transfers in phage functional and morphological diversification can be enhanced by the differences in host range between temperate and virulent phages. Both temperate and virulent phages require functions to regulate the lytic cycle, but the former need to establish additional genetic interactions with their hosts during lysogeny. This includes the regulation of lysis-lysogeny decision and the regulation of the induction of the lytic cycle in the prophage. These processes often require additional integration of the phage regulatory mechanisms in the cell’s genetic network (e.g., many phages are induced when their hosts’ SOS response is activated). This leads to a tighter host-parasite co-evolution, and potentially reduces the host range of the temperate phages, contributing to our observation that virulent phages have a relatively broader host range. This is also in agreement with the experimental observation that virulent coliphages isolated from the faeces of toddlers infect a broader range of gut strains (lysogens or otherwise) than temperate ones (Mathieu et al. 2020). The implications of our findings are that groups of temperate phages that might otherwise be sexually isolated (because they infect phylogenetically distant hosts) can exchange genes indirectly through exchanges with broader host range virulent phages. Conversely, it was observed that virulent-to-virulent genetic transfers are rare outside restricted taxonomic groups (Chopin et al. 2001; Lima-Mendez et al. 2008). Thus, temperate phages could also mediate exchanges between unrelated virulent phages. The genetic flux between virulent phages, and also between virulent and temperate phages, might then increase when they infect hosts with many prophages. This can accelerate the diversification of the gene repertoires of phage genomes and of lysogenic bacteria.

## Materials and Methods

### Data

We retrieved the complete genomes of 13513 bacteria and 2502 phages from NCBI non-redundant RefSeq database (ftp://ftp.ncbi.nlm.nih.gov/genomes/refseq/, last accessed in May 2019). 5 phage genomes were excluded because the annotation of the genomes lacked the gene identification, resulting in a dataset with 2487 phage genomes. The lifestyle of these phages was predicted using PHACTS v.0.3 (McNair et al. 2012). Predictions were considered as confident if the average probability score of the predicted lifestyle was at least two standard deviations (SD) away from the average probability score of the other lifestyle, which leads a precision rate in lifestyle identification of 99% (McNair et al. 2012). Using these criteria, we classified as confident 54% of the phages into 571 virulent and 780 temperate phages. Alternatively, we used BACPHLIP version 0.9.3 (Hockenberry and Wilke 2020) (default parameters) to predict the lifestyle of each phage, for almost all (> 99%) of the genomes. Data for the HGCF and LGCF phages was taken from (Mavrich and Hatfull 2017) by matching the NCBI identifiers.

We retrieved information (when available) on the phage hosts from the Genbank files of the phages or from the Virus-Host DB (Mihara et al. 2016) (https://www.genome.jp/virushostdb/). A few phages (~2%) have no identified host, since they were collected from environmental (soil, sea, etc.) or faeces samples. In this dataset, the phage hosts belong to 332 species and 145 genera. Most of these species (69%) have at least one complete genome sequenced and available in NCBI RefSeq. In the absence of a complete genome of the same species (31%, indicated as “manually annotated” in File S1), we used another genome from the same genus as a proxy for the species. Some analyses were repeated by excluding these phages with manually annotated (non-confident) host species, in order to provide a conservative control for these assignments.

Prophages (integrated temperate phages) were predicted using VirSorter v.1.0.3 (Roux et al. 2015) with the RefSeqABVir database in these bacterial genomes (corresponding to the host species/genus of the phages). The least confident predictions, i.e., categories 3 and 6, which may be prophage remnants or erroneous assignments, were excluded from the analyses. We also retrieved the phage family from the Genbank files of each phage. Most of them are Caudovirales (94%) and belong to the 3 main phage families, i.e., Siphoviridae (1291), Myoviridae (632) and Pododviridae (383). The complete dataset of phage genomes can be found in File S1.

### Protein similarity and weighted gene repertoire between bacteriophage genomes

We searched for sequence similarity between all proteins of all phages using mmseqs2 (Steinegger and Söding 2017) (Nature Biotechnology release, August 2017) with the sensitivity parameter set at 7.5. The results were converted to the blast format for analysis and we kept for analysis the hits respecting the following thresholds: e-value lower than 0.0001, at least 35% identity, and a coverage of at least 50% of the proteins. The hits were used to compute the bi-directional best hits between pair of phages, which were used to compute a score of gene repertoire relatedness weighted by sequence identity:

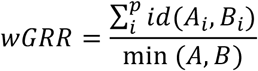

where A_i_ and B_i_ is the pair *i* of homologous proteins present in *A* and *B*, id(*A_i_*,*B_i_*) is the percent sequence identity of their alignment, and min(*A*,*B*) is the number of proteins of the genome encoding the fewest proteins (*A* or *B*). wGRR is the fraction of bi-directional best hits between two genomes weighted by the sequence identity of the homologs. It varies between zero (no bi-directional best hits) and one (all genes of the smallest genome have an identical homolog in the largest genome). wGRR integrates information on the frequency of homologs and sequence identity. Hence, when the smallest genome is 100 proteins long, a wGRR of 0.03 can result from three pairs of identical homologs or six pairs of homologs with 50% identity.

### Similarity networks, community detection and calculation of entropy

The phage network was built based on the filtered wGRR values, using the *networkx* and *graphviz* Python (v2.7) packages, and the *neato* algorithm. The Louvain community clusters (Blondel et al. 2008) were calculated using the *best_partition* function from the *community* package in Python (v2.7). For each cluster (considering only those with at least 3 nodes), we first calculated their total number of nodes, and then the number of nodes corresponding to each category: nodes in either one of two lifestyles (temperate or virulent), nodes with a given phage family and nodes with a given described bacterial host phyla. The Shannon entropy of a cluster (X) with nodes classed according to a given variable that takes *t* different values (*y_1,X_*… *y_t,X_*) was calculated as:

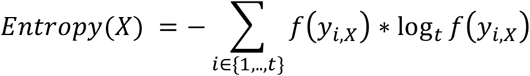

Where *f*(*y_i,X_*) is the relative frequency of the nodes classed *i* in the cluster. As an example, consider a cluster composed of 20 nodes, where 10 of them have a temperate lifestyle and 10 have a virulent lifestyle. Because there are two labels (temperate or virulent), *t* = 2. If the frequencies of the two types of phages were identical, close to the expectation under a random distribution, the two frequencies would be 0.5 and the entropy would amount to 1, which is the maximum and corresponds to a highly heterogeneous cluster. When all phages are from one single type, the entropy is equal to 0, corresponding to an homogeneous cluster. To be able to compare each trait to the expectation of random cluster compositions, we randomly re-assigned the labels (host phyla, phage family and lifestyle) to the phages and computed the Shannon entropy as described above. The results (shown as boxplots in the figures) summarise the distribution of the 100 random experiments.

### Identification of pairs of phage genomes with recombinant genes

In order to separate events of recombination and from homology due to shared ancestry, we used the relationship between the fraction of homologous proteins (over the average number of proteins for each phage pair) and the fraction of homologous proteins with very high identity (at least 80% identity, also over the average number of proteins for each phage pair). We used the *ols* function from the *statsmodels* package in Python (v2.7) to generate a linear model describing the expectation of ancestry between phage genomes. To characterize common ancestry as robustly as possible, we restricted the dataset to temperate-virulent pairs of phages, as this dataset is expected to contain the lowest frequency of recombination events. Further, and in order to reduce the influence of the outliers associated with recombination (and thus maximize the fit of the linear model to cases of common ancestry), we analysed only the pairs of phage with at least 50% of the proteins in their (average) genomes classified as homologous (meaning at least 35% identity and 50% coverage) and a minimum of one highly similar homologous protein (at least 80% identity). Using this dataset, we applied residual analysis (with the *fit* and *outlier_test* function from *statsmodels*) to identify the first significant residual value. This was defined as the least distant residual from the linear model that is classed as outside the confidence interval, with an adjusted pvalue<0.05 using the Benjamini/Hochberg method – *fdr_bh*. The value of this residual is assumed as the significance threshold beyond which a residual is classified as a putative recombination event (since it significantly departs from the expectation of common ancestry defined by the linear model). Thus, the linear model and the residual threshold are applied to the entire range of the data across the three datasets (temperate-virulent, temperate-temperate and virulent-virulent pairs). The pairs of phages whose residuals are larger than the minimum significant residual threshold (and whose wGRR are below 0.5 to avoid high similarity caused by recent divergence of the lineages) are classed as pairs of putatively recombined phages. The recombinant genes are identified as the proteins with at least 80% identity between the pairs of phages showing evidence of recombination.

### Functional annotation of proteins

We used HMMER v3.1b2 (Eddy 2011) (default parameters) to search for genes matching the prokaryotic Virus Orthologous Groups (pVOGS) (Version May 2016 (Grazziotin et al. 2017)) database of hmm profiles (filters used were e-value<1e-5 and profile coverage>60%). Only pVOGs with a viral quotient (VQ) above 0.8 were used (7751 out of 9518 pVOG profiles in total). The pVOG profiles were classed into seven functional categories: a) structure, b) lysis, c) packaging, maturation/Assembly and DNA injection, d) DNA metabolism, recombination and regulation, e) others and f) unknown by two approaches. First a profile-profile comparison was done using the HHsuite 2.0.9 (Söding 2005) with phage-specific profiles from the PFAM (El-Gebali et al. 2019) and TIGRFAM (Haft et al. 2003) database (taken from (Fouts 2006)). Applying a threshold of p-value <10^−5^ resulted in 711 profiles that cluster in 261 groups using the Louvain algorithm. The annotations of the PFAM and TIGFRAM profiles were used to assign one of the functional categories to a group. The remaining profiles were manually assigned considering the piled-up annotations of the pVOGs (File S3). The identification of bacterial functions was performed based the EggNOG database of hmm profiles (bactNOG Version (Huerta-Cepas et al. 2019)). In order to minimize the number of bactNOG entries that derive from prophages in bacterial genomes, we used hhsearch (HHSuite version 3.2 (Steinegger et al. 2019)) to remove from bactNOG the profiles matching pVOG profiles. The bactNOG profiles with matches in pVOGs with VQ above 0.8 (p-value <0.0001) were discarded. This resulted in a reduction of 34% of the bactNOG profiles’ dataset. The remaining 135814 bactNOG profiles were used to class the dataset of putative recombinant genes (filters used were e-value<1e-5 and profile coverage>60%) in broad functional categories.

### Detection of phage-like recombinases in phage and bacterial genomes

The families of recombinases of phage were described in (Lopes et al. 2010), for which we built profiles or recovered them from PFAM. To build the profiles, we retrieved the homologs given in the reference above and aligned them using default options with MAFFT (v7.407) (Katoh and Standley 2013). The alignments were used to build the profiles using hmmbuild from HMMER (default options). Our novel profiles are given in supplementary material (File S6). We used HMMER to search for homologs of Sak (PF04098, --cut_ga), Sak4 (Sak4 from phage T7, minimum score = 20), Erf (PF04404, --cut_ga), RecT (PF03837, --cut_ga) and gp2.5 (gp2.5 from phage T7, minimum score = 20). An additional profile, UvsX, also matching RecA (PF00154, --cut_ga) was searched only in phage genomes. Hosts were retrieved from the GenBank file of each phage, and all the genomes in the database belonging to that host’s species were used for the calculation. E.g., if the host species of a given phage is described as *E. coli*, we calculated the mean number of recombinases in all *E. coli* genomes. The values shown in the table are the mean of these means for phages with recombinant genes and the remaining phages. If recombinases were found within the coordinates of a prophage, they were associated with prophage regions. Note that a particular phage host can be included multiple times in the distribution, if subsequent phages have a similar host, and can even result in the host being included in the dataset of recombinant and non-recombinant phages simultaneously. Although this results in repeated sampling, it represents the likelihood that hosts of recombinant or non-recombinant phages encode for recombinases. Moreover, this is unbiased for either class of phages, since the hosts of both recombinant and non-recombinant phages are subject to this repeated sampling process.

### Detection of NHEJ in bacterial genomes

We used HMMER to search for homologs of the Ku protein (PF02735.16) in the dataset of bacterial genomes, using the -cut_ga parameter. We did not require the presence of a neighboring ligase to consider the system complete, because such a ligase is often absent (Bernheim et al. 2019). The set of bacterial genomes assigned as host of a given phage was asserted as described in the section above. The fraction of hosts that encode for NHEJ was calculated as the number of genomes that where at least one homolog of Ku was found, divided by the total number of genomes considered for a given phage. The use of the presence/absence of Ku as the key variable, and not the number of Ku genes, is due to three reasons: 1) the numbers were always low (most often 0 or 1), 2) some genomes encoding multiple NHEJ systems express them during different times, 3) some bacteria seem to encode heterodimeric Ku where two genes are necessary for the process (Bertrand et al. 2019).

### Detection of IS in phage genomes

We used HMMER to search for the profiles of transposases contained in the ISEScan tool (Xie and Tang 2017) in phage genomes. We retained all hits with an e-value of at most 1e-5, and a coverage of at least 60%, for at least one of the profiles in the collection.

### Calculation of the patristic distance between bacterial hosts

We used the 16S rRNA of the bacterial genomes identified in the RefSeq annotations, corresponding to the species level of the identified phage hosts. We selected the first entry of each genome and aligned them using the secondary structure models with the program SSU_Align version 0.1.1 (Nawrocki et al. 2009). Poorly aligned positions were eliminated with the ssu-mask. The alignment was trimmed with trimAl version 1.2 (Capella-Gutiérrez et al. 2009) using the option –noallgaps to delete only the gap proteins but not the regions that are poorly conserved. The 16S rRNA phylogenetic tree was inferred using maximum likelihood with IQTREE version 1.6.5 (Nguyen et al. 2015) under the best-fit model automatically selected by ModelFinder (Kalyaanamoorthy et al. 2017), and with the options –wbtl (to conserve all optimal trees and their branch lengths) and –bb 1000 to run the ultrafast bootstrap option with 1000 replicates. The patristic distances amongst the taxa in the 16S trees were calculated from the tree (weighted by the edge distances) with the *dendropy* package in Python (Sukumaran and Holder 2010), using the functions *phylogenetic_distance_matrix* and *patristic_distance*, with the default parameters.

### Calculation of the genetic distance between bacterial hosts

We calculated the trinucleotide composition, or 3-mer genomic signature, of each bacterial genome from the phage host species, using the relative abundance value of each of the 64 possible trinucleotides, as in (Suzuki et al. 2010). This is defined as the observed trinucleotide frequency divided by the expected trinucleotide frequency (the product of the mononucleotide frequencies), or

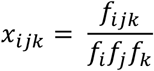

where *f_i_*, *f_j_* and *f_k_* represent the frequency of the nucleotides *i*, *j* and *k*, respectively (with *i*, *j*, *k* ∈ A, C, T, G). This allows to quantify the deviation of the observed frequency of trinucleotides to the one expected given the nucleotide composition of the genomes, which are known to differ between phages and bacteria (Rocha and Danchin 2002). The genetic distance between two hosts was then calculated using the average absolute difference between the 3-mer genomic composition of each, as

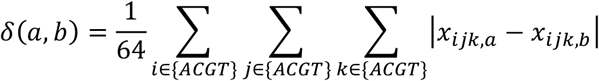

where *x_ijk,a_* and *x_ijk,b_* are the relative abundances of the trinucleotide *ijk* in each of the bacterial hosts. Note that multiple genomes might be available for each host. To use all the available information when multiple genomes are available for a species, we used the grand mean of all pairwise comparisons. As an example, if one host is *Escherichia coli* and the another is *Staphylococcus aureus*, we calculated the differences in genomic signature between each pairwise combinations of all genomes of *Escherichia coli* and all genomes of *Staphylococcus aureus*. We then calculate the average of all the pairwise calculations of the genetic distance as *δ_T_*, with

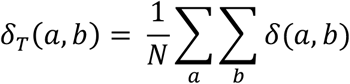

where *a* and *b* are individual strain genomes of the first and second host, respectively, and *N* is the total pairwise calculations between the strains of the first and second hosts.

## ACKNOWLEDGEMENTS

We acknowledge support by the PRESTIGE programme (PRESTIGE-2017-1-0012), the ANR grants (ANR-16-CE16-0029), the Fondation pour la Recherche Médicale (Equipe FRM EQU201903007835), and the Laboratoire d’Excellence IBEID (ANR-10-LABX-62-IBEID). We thank Mireille Ansaldi, Marta Lourenço and the members of the Microbial Evolutionary Genomics laboratory by discussions and comments on earlier versions of this manuscript. This work used the computational and storage services (TARS cluster) provided by the IT department at Institut Pasteur, Paris.

## DATA AVAILABILITY

All bacterial and bacteriophage sequences were retrieved from RefSeq, and are publicly available. The in-house scripts developed for the analysis and for constructing the figures are available upon request.

