## Supplementary information file for "Causes and consequences of bacteriophage diversification via genetic exchanges across lifestyles and bacterial taxa"

**This PDF file includes:**

Figures S1 to S20
Tables S1 to S7
Legends for Datasets S1 to S6

**Other supplementary materials for this manuscript include the following:**

Datasets S1 to S6

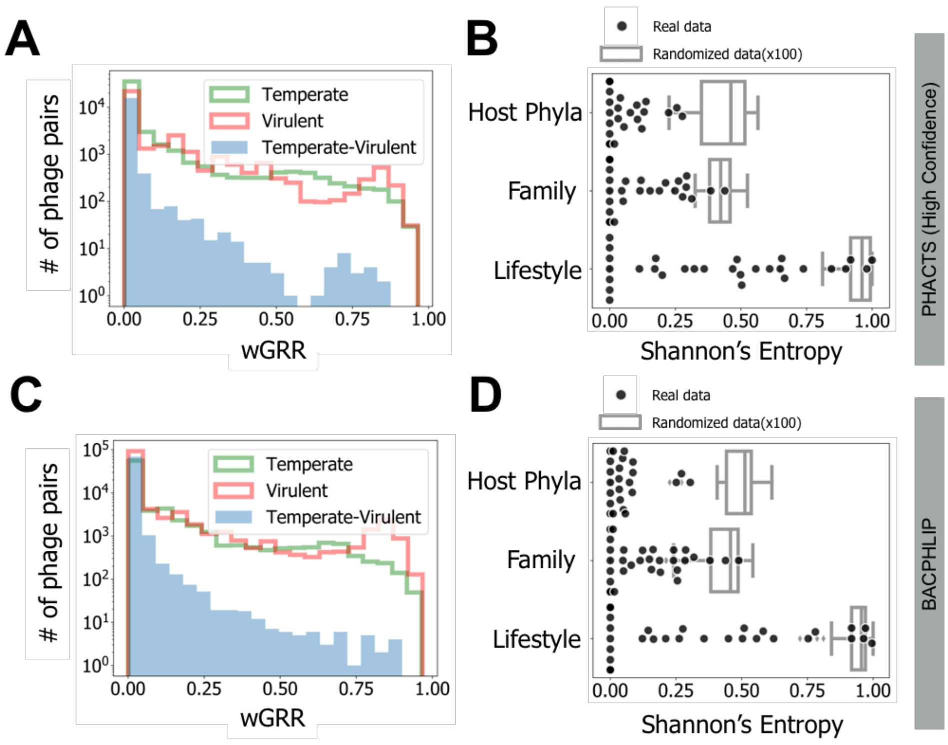


**Fig. S1. Distributions of homology between phages and heterogeneity of the clusters of the corresponding wGRR network using alternative phage lifestyle classifications.** In A and B, only the phages' lifestyles confidently assigned by PHACTS. In C and D), phages lifestyles assigned by BACPHLIP. **A and C)** Histograms of the wGRR values (with wGRR>0). **B)** Shannon's Entropy values for each cluster identified with the Louvain community detection from the wGRR matrix, for the three phage traits (N=29 for each trait, one per cluster). Boxplots represent the distribution of the concatenation of 100 repetitions of randomized cluster label re-assignments (N=2900 for each trait). All distributions are significantly less heterogeneous than their random counterparts (p<1e-22 for Host Phyla, p<1e-17 for Family and p<1e-14 for Lifestyle, 2-sample Kolmogorov Smirnoff test), and the real clusters are significantly more heterogeneous in phage lifestyle (Tukey Honest Significant Difference (HSD) Test p<0.001 for Lifestyle versus Host Phyla or Family, p=0.5 for Host Phyla versus Family). **D)** Shannon's Entropy values for each cluster identified with the Louvain community detection from the wGRR matrix, for the three phage traits (N=53 for each trait, one per cluster). Boxplots represent the distribution of the concatenation of 100 repetitions of randomized cluster label re-assignments (N=5300 for each trait). All distributions are significantly less heterogeneous than their random counterparts (p<1e-45 for Host Phyla, p<1e-21 for Family and p<1e-20 for Lifestyle, 2-sample Kolmogorov Smirnoff test), and the real clusters are significantly more heterogeneous in phage lifestyle (Tukey Honest Significant Difference (HSD) Test p<0.001 for Lifestyle versus Host Phyla or Family, p=0.03 for Host Phyla versus Family).

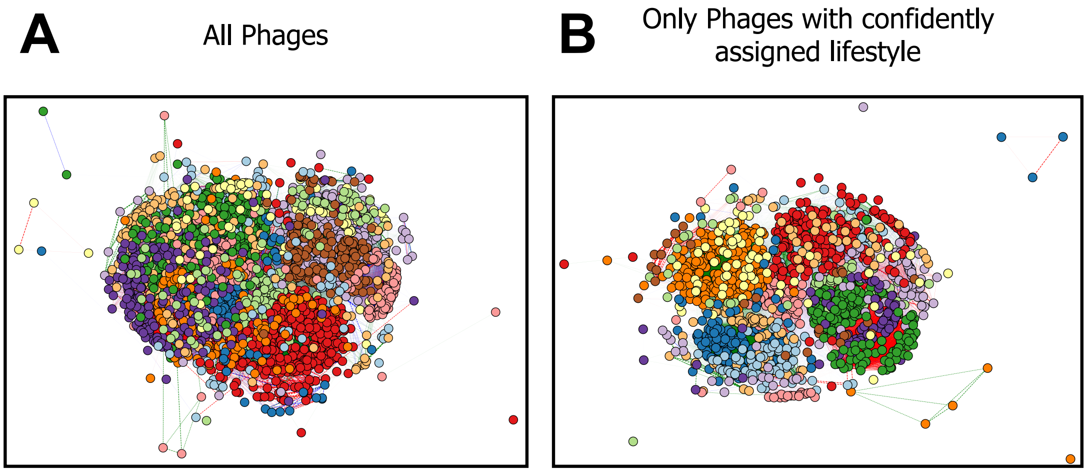


**Fig. S2. Phage network based on the complete wGRR matrix. A)** All phages considered for the network. **B)** Only phages with a confidently assigned lifestyle were considered for the network. [Both panels] Each node correspond to a phage, and the edges correspond to a homology link between two phages (wGRR >0). Node colors correspond to the clusters identified by the Louvain community detection method. Edge colors correspond to the connection between temperate-temperate (green), temperate-virulent (blue) or virulent-virulent (red) phages.

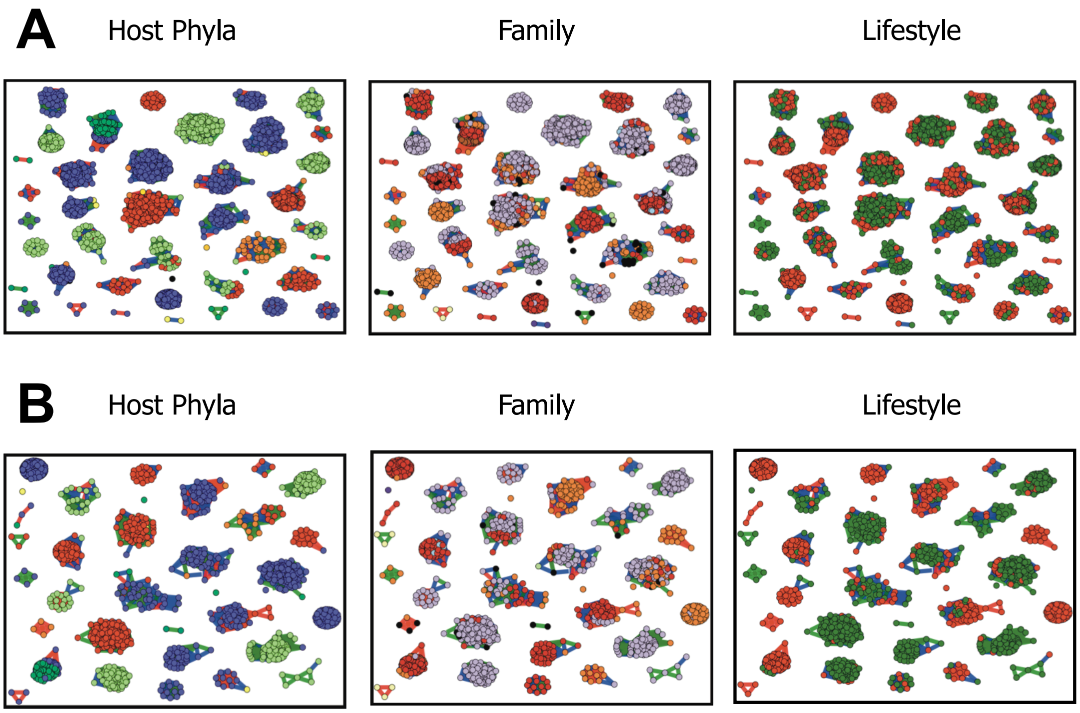


**Fig. S3. Individual clusters identified with the Louvain community detection method represented as separate components. A)** Clusters from the network considering all phages. In the leftmost panel, node colors represent the different phyla of the phage's bacterial hosts, in the central panel node colors represent different phage families and in the rightmost panel colors represent the lifestyle of phages (temperate in green, virulent in red). In all panels edge colors correspond to the connection between temperate-temperate (green), temperate-virulent (blue) or virulent-virulent (red) phages. **B)** Same as in **A**, but only considering phages with a confidently assigned lifestyle in PHACTS.

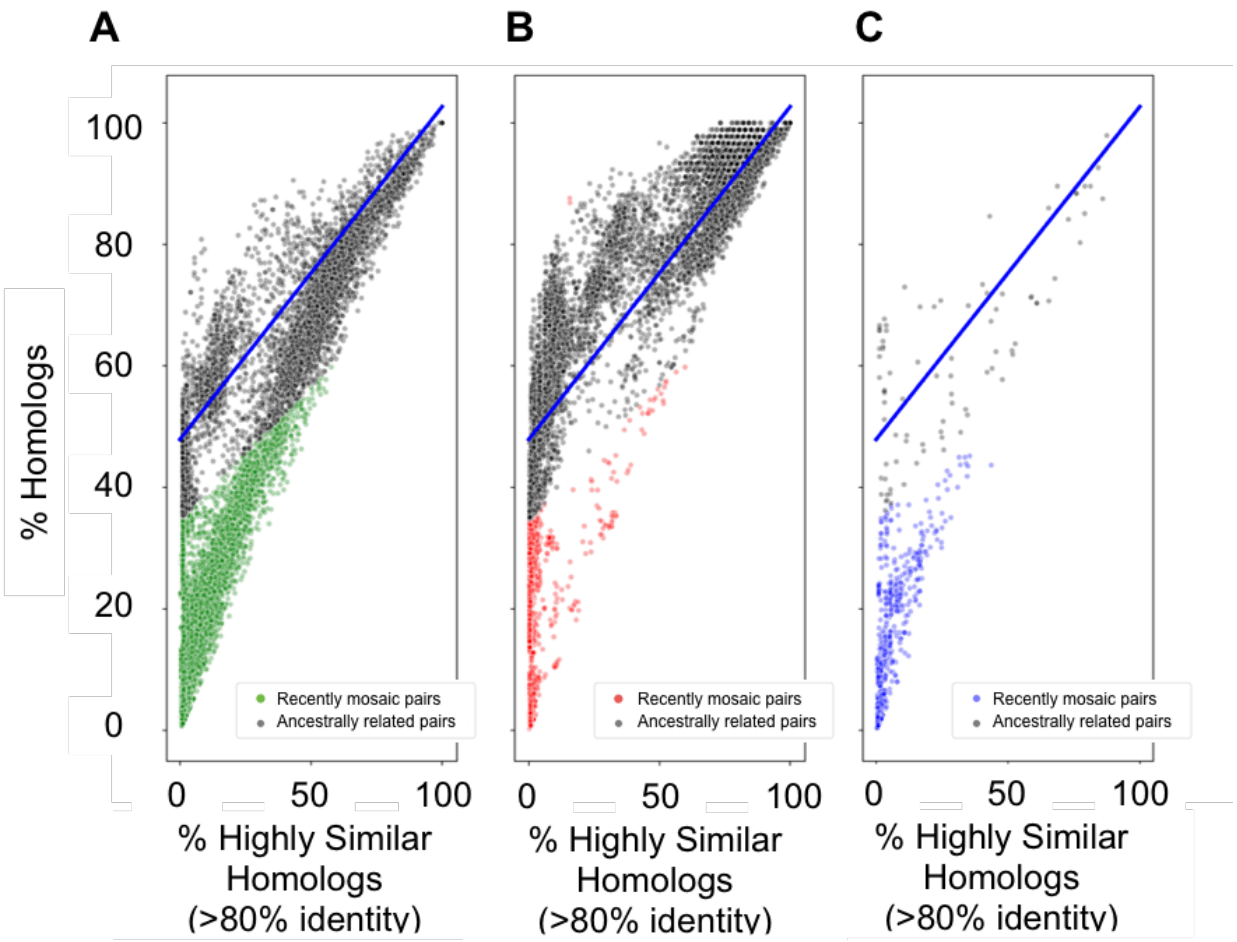


**Fig. S4. Identification of putative recently mosaic phage pairs with lifestyle assigned with BACPHLIP. A-C)** Scatterplots of pairs of phage genomes with recently mosaic phage pairs indicated as colored points (otherwise grey). The linear regression model (blue line) was inferred for the virulent-virulent dataset in B and applied to all datasets. **A)** Pairs of temperate phages. **B)** Pairs of virulent phages. **C)** Pairs of temperate-virulent phages.

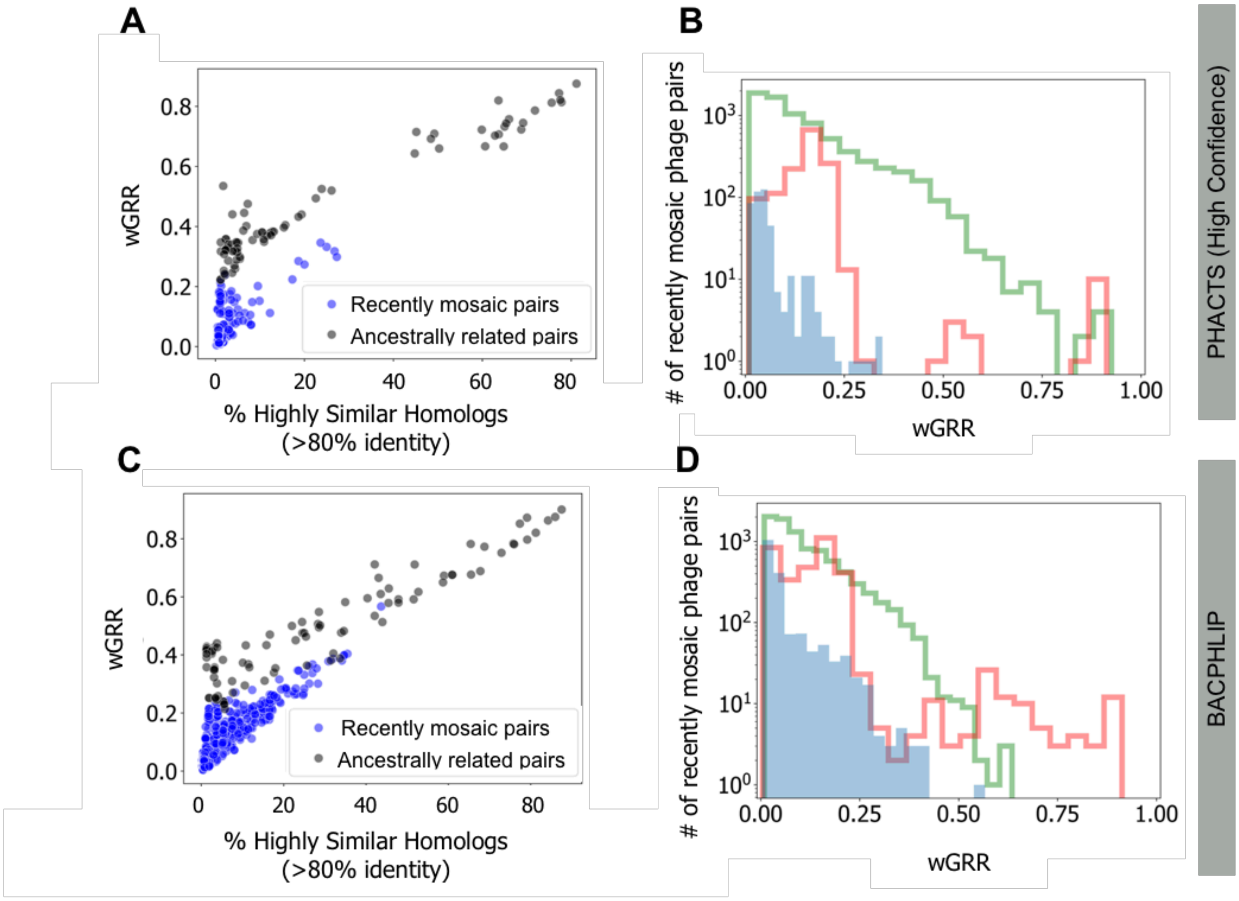


**Fig. S5. wGRR distributions of putative recently mosaic phage pairs, whose phages a lifestyle assigned with alternative approaches.** In A and B, only the phages' lifestyles confidently assigned by PHACTS. In C and D), phages lifestyles assigned by BACPHLIP. **A and C)** Scatterplot of pairs of temperate-virulent phages in terms of wGRR and the fraction of high sequence-identity homologous genes. **B and D)** Histogram of wGRR values for the subset of recombinant phage pairs. For C and D (BACPHLIP), data originates from Fig. S4.

**A**

| | Temperate | Virulent |
|---|---|---|
| **LGCF** | 227 | 510 |
| **HGCF** | 249 | 34 |
| **Unclassified** | 821 | 546 |

**B**

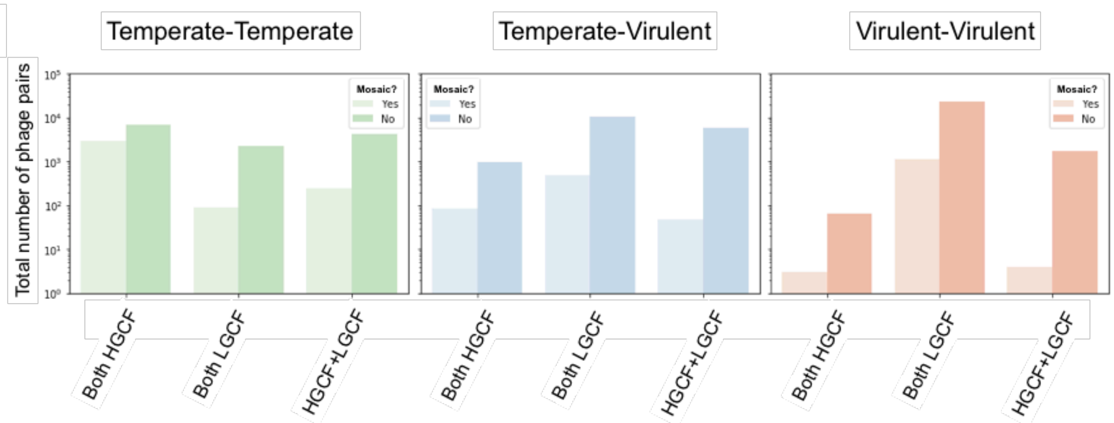


**Fig. S6. Gene content flux mode identified for the recently mosaic phage pairs. A)** Total number of phages in the dataset classified as HGCF or LGCF (last row indicates phages that have either an unknown/mixed classification on the Maverick dataset, or phages that are missing from that dataset and are thus are unclassified regarding their mode of gene content flux). B) Total number of phage pairs (identified as recently mosaic or otherwise) with each combination of gene content flux mode (i.e., both phages have similar GCF mode, or each phage in the pair has a different GCF mode). Each panel represents a different lifestyle combination between the pairs of phages.

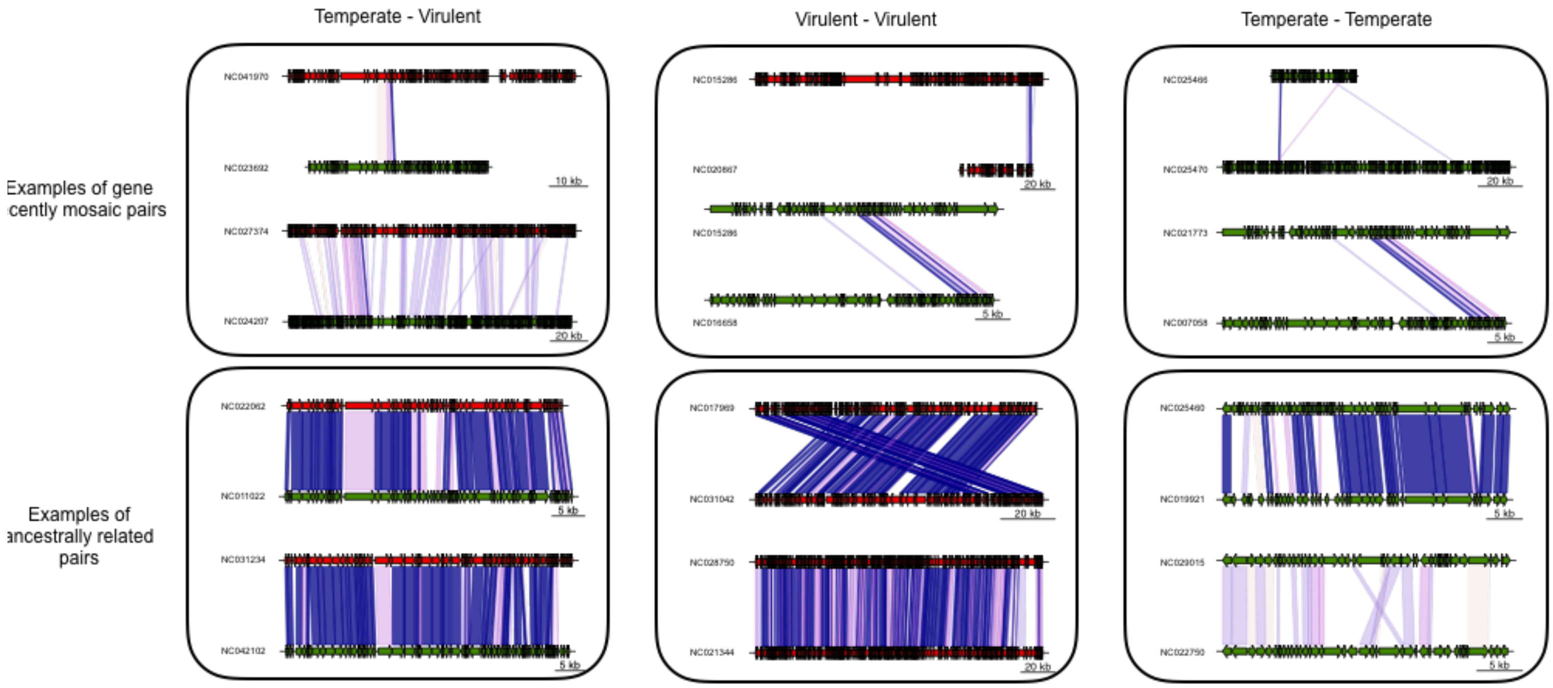


**Fig. S7. Examples of pairs of phages classified as recently mosaic or remaining (ancestrally related).** Color codes for genes: virulent (red), temperate (green), virulent-temperate (blue). Colors in the blocks linking the phages indicate sequence similarity between homologs.

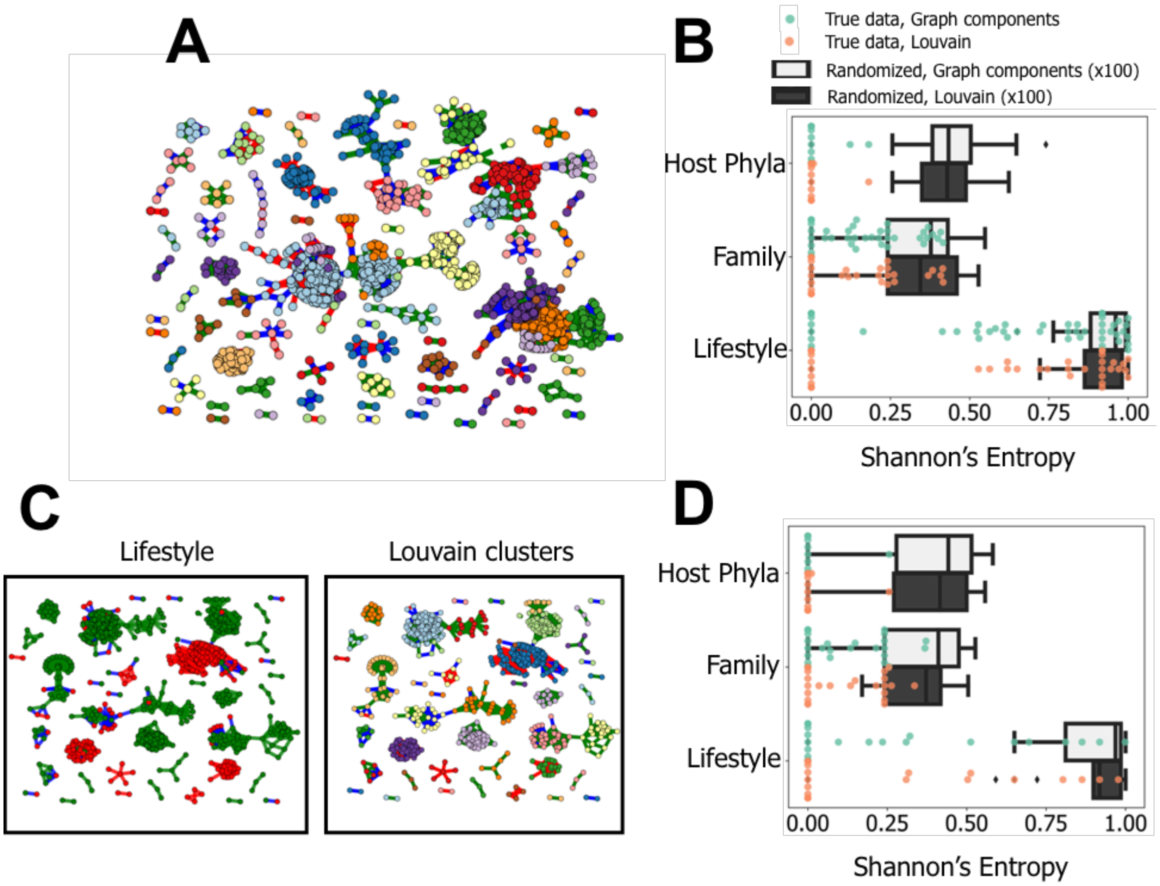


**Fig. S8. Network of recently mosaic phages and quantification of cluster heterogeneity. A)** Same network of recombinant phages as the one in Figure 3D in the main text (considering all phages), but with each node colored by its presence in the clusters identified with the Louvain community detection method. Edge colors correspond to the connection between temperate-temperate (green), temperate-virulent (blue) or virulent-virulent (red) phages. **B)** Shannon's Entropy values for each of the three phage traits in each of the clusters identified either from directly from the graph ("graph components", shown in turquoise points and white boxplots, N=38 and 3800, respectively) or with the Louvain community detection (panel **A**, shown in orange points and black boxplots, N=56 and 5600, respectively), Boxplots represent the distribution of the concatenation of 100 repetitions of randomized cluster label re-assignments. All distributions are significantly less heterogeneous than their random counterparts (in "graph components", $p<1e-32$ for Host Phyla, $p<1e-10$ for Family and $p<1e-04$ for Lifestyle; in Louvain, $p<1e-46$ for Host Phyla, $p<1e-15$ for Family and $p<1e-07$ for Lifestyle, 2-sample Kolmogorov Smirnoff test), and the real clusters are significantly more heterogeneous in phage lifestyle (in "graph components", $p<0.001$ for Lifestyle versus Host Phyla or Family, $p=0.12$ for Host Phyla versus Family; in Louvain, $p<0.001$ for Lifestyle versus Host Phyla or Family, $p=0.03$ for Host Phyla versus Family in Louvain, Tukey Honest Significant Difference Test). **C)** Same networks as in Figure 3C (rightmost panel) or panel **A**, but considering only phages with a confidently assigned lifestyle. **D)** Shannon's Entropy values for each of the three phage traits in each of the clusters from the network with only phages with a confidently assigned lifestyle, identified either from directly from the graph ("graph components", shown in turquoise points and white boxplots, N=23 and 2300, respectively) or with the Louvain community detection (panel **A**, shown in orange points and black boxplots, N=29 and 2900, respectively), Boxplots represent the distribution of the concatenation of 100 repetitions of randomized cluster label re-assignments. All distributions are significantly less heterogeneous than their random counterparts (in “graph components”, $p<1e-17$ for Host Phyla, $p<1e-07$ for Family and $p<1e-11$ for Lifestyle; in Louvain, $p<1e-22$ for Host Phyla, $p<1e-09$ for Family and $p<1e-10$ for Lifestyle, 2-sample Kolmogorov Smirnoff test), and the real clusters are significantly more heterogeneous in phage lifestyle (in “graph components”, $p<0.001$ for Lifestyle versus Host Phyla or Family, $p=0.12$ for Host Phyla versus Family; in Louvain, $p<0.001$ for Lifestyle versus Host Phyla or Family, $p=0.03$ for Host Phyla versus Family in Louvain, Tukey Honest Significant Difference Test).

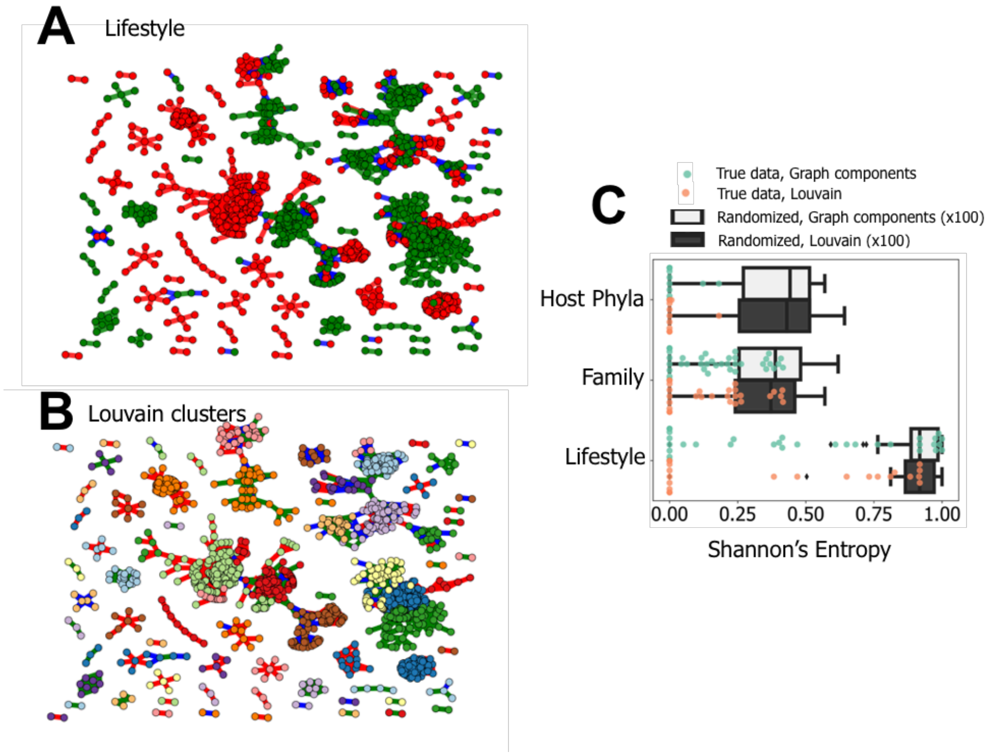


**Fig. S9. Network of recently mosaic phages whose lifestyle was inferred using BACPHLIP, and quantification of its clusters' heterogeneity.** Recombination data originates from Fig S4. **A)** Network of recombinant phages. Each node represents a phage genome and each edge a relationship of genetic similarity (with wGRR below 0.5). Edge colors correspond to the connections between temperate-temperate (green), temperate-virulent (blue) or virulent-virulent (red) phages. **B)** Same as in A, but with each node colored by its presence in the clusters identified with the Louvain community detection method. **C)** Shannon's Entropy values for each of the three phage traits in each of the clusters from the network with only phages with a confidently assigned lifestyle, identified either from directly from the graph ("graph components", shown in turquoise points and white boxplots, N=36 and 3600, respectively) or with the Louvain community detection (panel A, shown in orange points and black boxplots, N=53 and 5300, respectively), Boxplots represent the distribution of the concatenation of 100 repetitions of randomized cluster label re-assignments. All distributions are significantly less heterogeneous than their random counterparts (in "graph components", p<1e-26 for Host Phyla, p<1e-09 for Family and p<1e-14 for Lifestyle; in Louvain, p<1e-45 for Host Phyla, p<1e-21 for Family and p<1e-20 for Lifestyle, 2-sample Kolmogorov Smirnoff test), and the real clusters are significantly more heterogeneous in phage lifestyle (in "graph components", p<0.001 for Lifestyle versus Host Phyla, p=0.06 for Lifestyle vs Family, p=0.03 for Host Phyla versus Family; in Louvain, p<0.001 for Lifestyle versus Host Phyla or Family, p=0.03 for Host Phyla versus Family in Louvain, Tukey Honest Significant Difference Test).

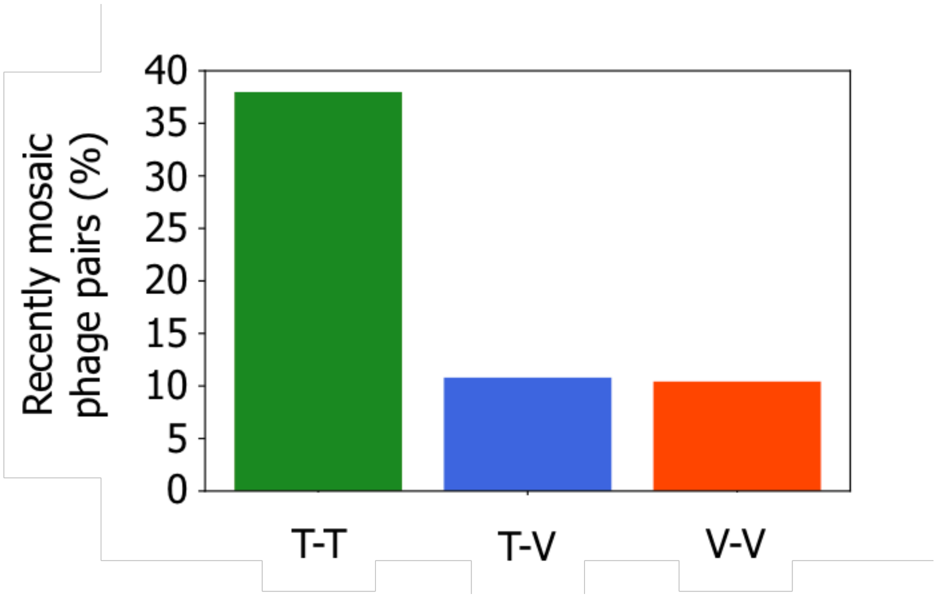


**Fig. S10. Quantification of impact of phage lifestyle on the frequency of recently mosaic phage pairs.** Mosaic plot of the association between the lifestyle of the pairs of phages (with wGRR>0.05) and the frequency with which a genetic transfer is identified to occur between them (p<0.001 for association with lifestyle, Chi2 test).

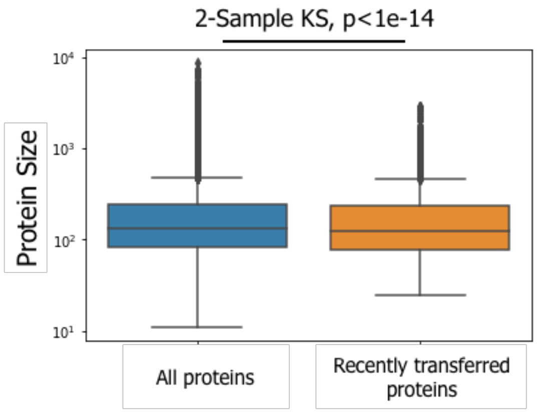


**Fig. S11. Size distributions of proteins in the phage dataset.** All proteins dataset refers the entire protein dataset of the phage collection (N=260145). Recently transferred proteins dataset refers to the homologous proteins with at least 80% identity found in the pairs of phages that were inferred as recently mosaic (N=21644).

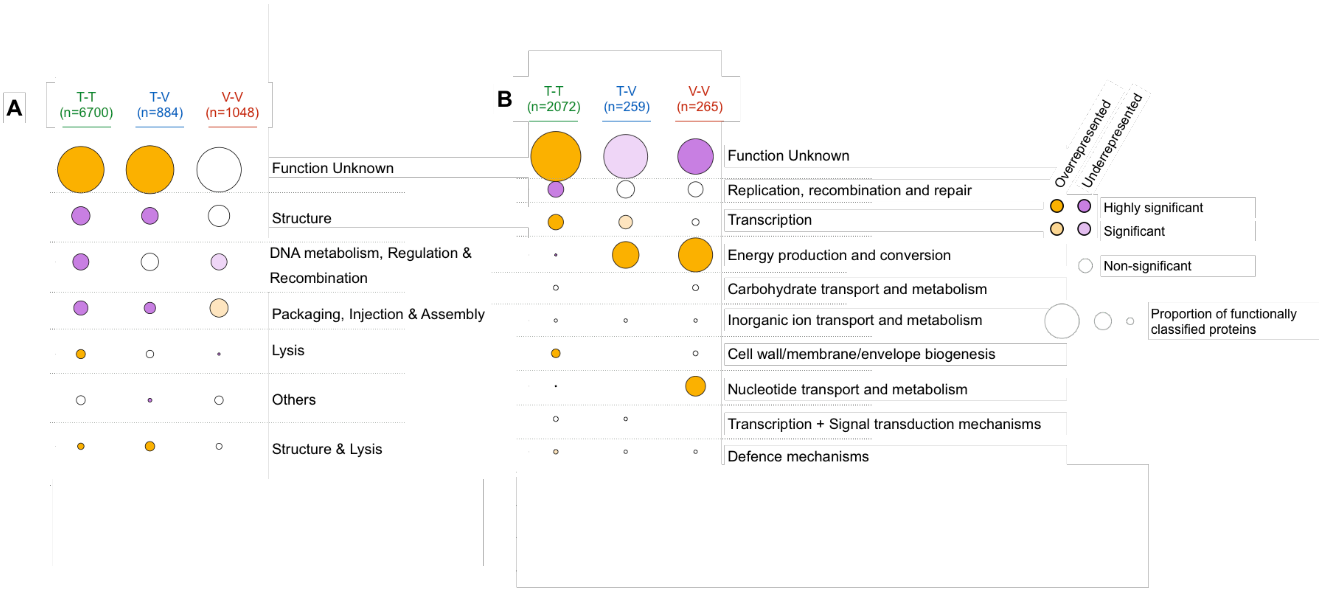


**Fig. S12. Functional classification of genes involved in genetic transfers between distantly related phages, with the phage lifestyles' inferred with BACPHLIP. A)** Phage-like functional classification using the pVOG database. **B**) Bacteria-like functional classification using the bactNOG database. In both panels, the size of each circle corresponds to the proportion of genes with a given function, for each lifestyle combination of phage pairs. The total number of genes with an assigned function is shown at the top, for each lifestyle combination. Enrichment (orange) or depletion (purple) of each function (per lifestyle combination) is assessed relatively to classification of all phage proteins with pVOG (in A) or bactNOG (in B). In both cases, the color intensity corresponds to the statistical significance of the classification (Fisher-exact test adjusted for multiple comparisons with the Benjamini-Hochberg method).

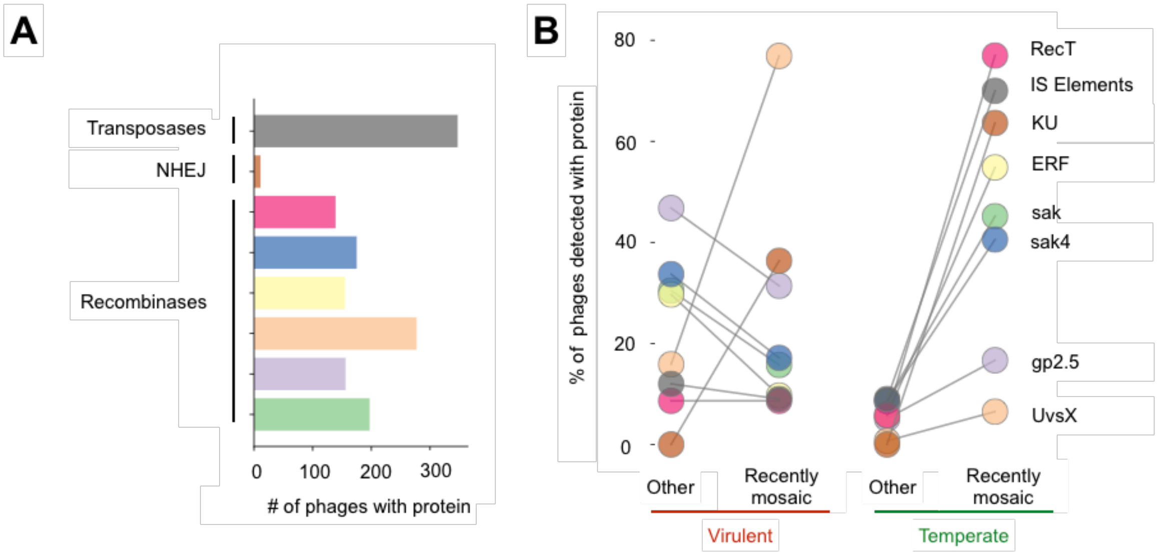


**Fig S13. Putative mechanisms involved in genetic transfers and their relative frequency in recently mosaic phages, with the phage lifestyles' inferred with BACPHLIP. A)** The total number of phages with at least one homologous gene for each of the proteins' types. **B)** Proportion of genomes with homologous genes for each type of proteins analysed in recently mosaic phages and the others. Colors of the bars and circles correspond to the different types of proteins analysed.

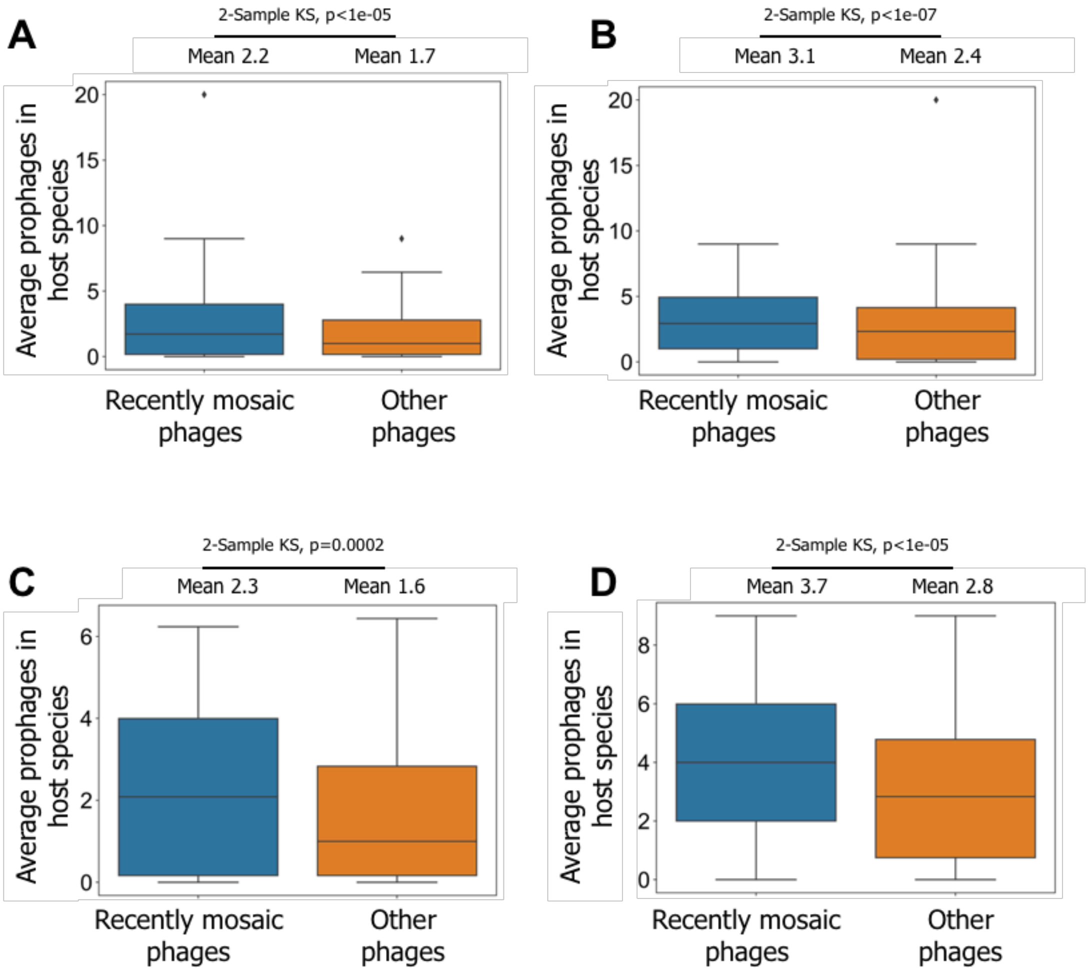


**Fig. S14. Distributions of the number of prophages in the phages' bacterial host species.** Boxplots represent the distributions of the average number of prophages in all the genomes of each phage bacterial host species, for recently mosaic (blue) or remaining (orange) phages. **A)** All temperate phages. **B)** All virulent phages. **C**) Only phages with a lifestyle confidently assigned as temperate. **D)** Only virulent phages with a lifestyle confidently assigned as virulent. Mean of the distribution and statistical significance (2 sample Kolmogorov-Smirnov) shown above each panel.

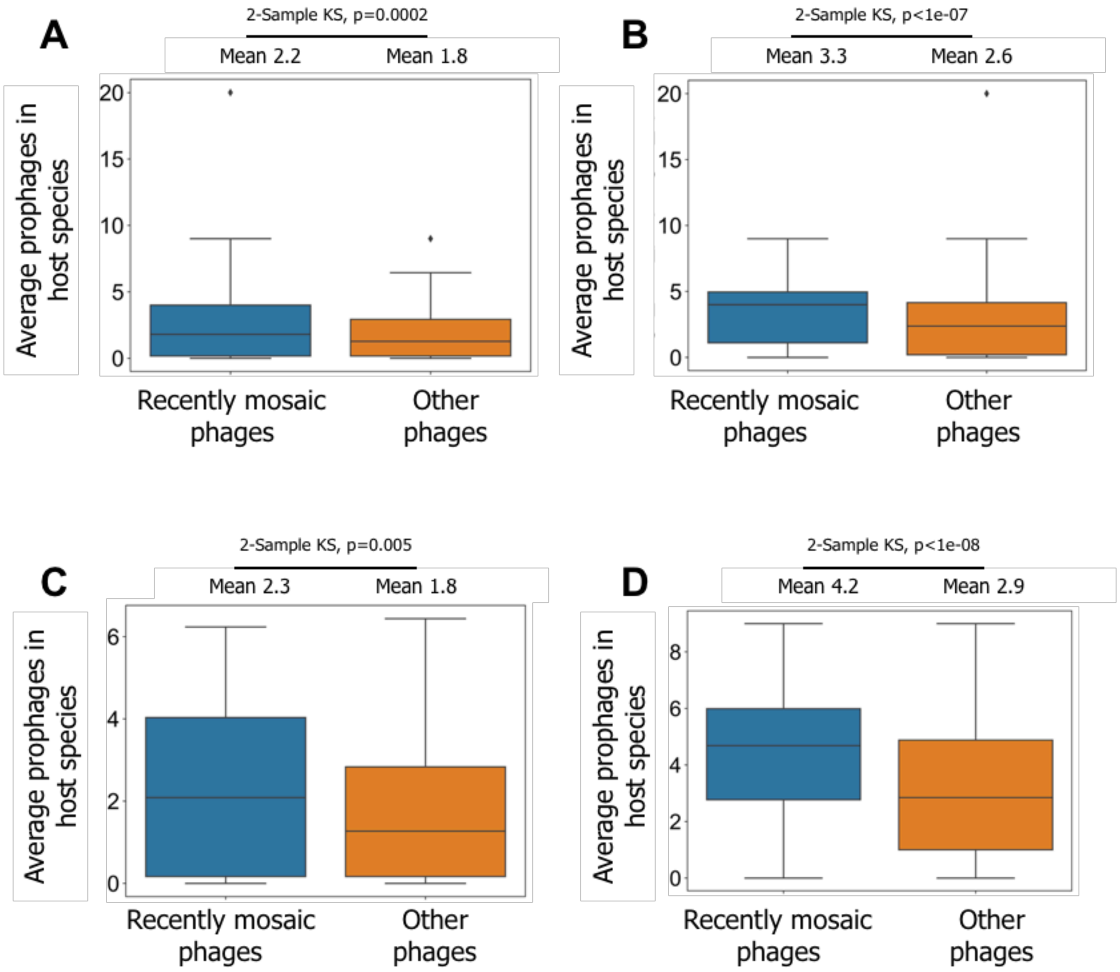


**Fig. S15. Distributions of the number of prophages in the phages' bacterial host species, considering only phages with a known annotated host species.** Same analysis as in Figure S7, but only considering phages with known annotated host species. **A)** All temperate phages. **B)** All virulent phages. **C**) Only phages with a lifestyle confidently assigned as temperate. **D)** Only virulent phages with a lifestyle confidently assigned as virulent. Mean of the distribution and statistical significance (2 sample Kolmogorov-Smirnov) shown above each panel.

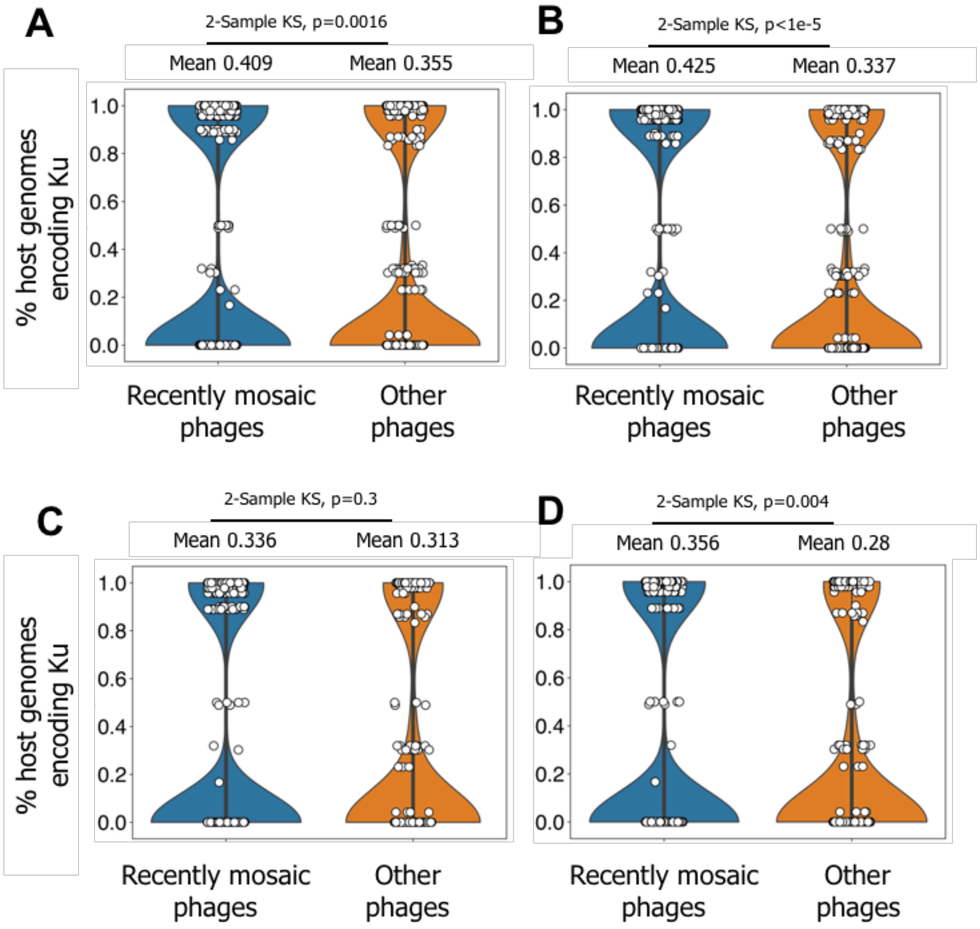


**Fig. S16. Fraction of the phages' bacterial host species with the Ku protein from the non-homologous end joining pathway. A)** All phages. **B)** All phages with a confidently assigned host species. **C)** Only phages with a confidently assigned lifestyle. **D)** Only phages with a confidently assigned lifestyle and a confidently assigned host species. Mean of the distribution and statistical significance (2 sample Kolmogorov-Smirnov) shown above each panel.

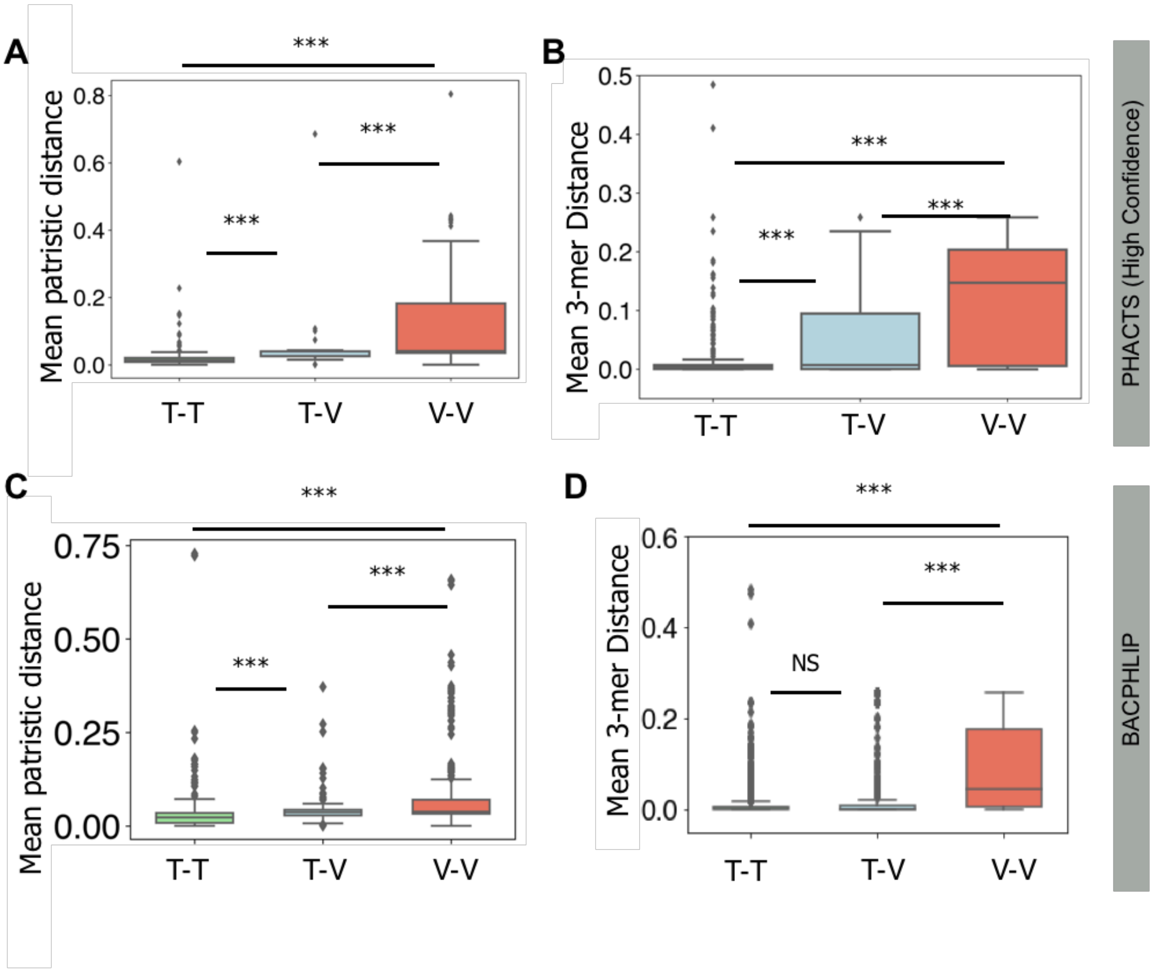


**Fig. S17. Distributions of the patristic distance and 3-mer genomic distance between the host species of recently mosaic phages, considering alternative phage lifestyle assignemnt approaches.** A and B, PHACTS using only phages with a confidently assigned lifestyle. C and D, lifestyle assignment with BACPHLIP. **A and C)** Boxplots of the non-null patristic distances (computed from the 16S rDNA gene tree) between the hosts' species of each recombinant phage pair. **B and D)** Distributions of the differences in 3-mer genomic signatures between the hosts's species of each recombining phage pair. *** p=0.001, Tukey HSD for all pairs. T-T: temperate-temperate, T-V: temperate-virulent, V-V: virulent-virulent. For C and D, data originates from Fig S4.

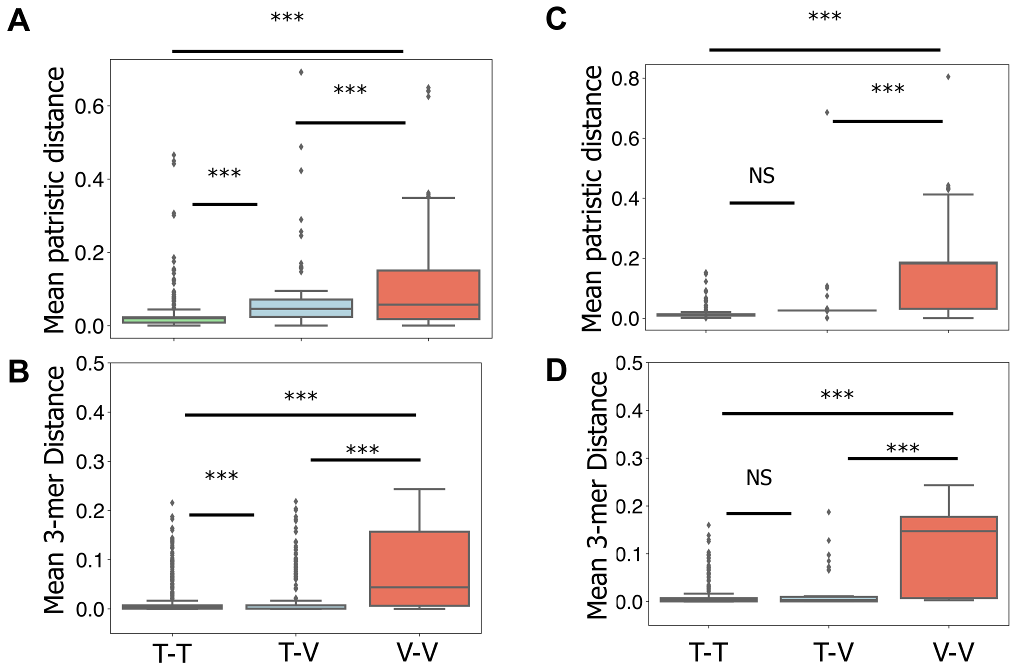


**Fig. S18. Distributions of the patristic distance and 3-mer genomic distance between the host species of recently mosaic phages, considering only phages with a known annotated host species.** Similar analysis to that in Figure 4 in the main text and Figure S15. **A)** and **B)** Phages with both confidently and non-confidently assigned lifestyles, but considering only those phages that have a known annotated host species. **C)** and **D)** Only phages with a confidently assigned lifestyle and with a known annotated host species. *** p=0.001, NS meaning non-significant (p>0.05), Tukey HSD for all pairs. T-T: temperate-temperate, T-V: temperate-virulent, V-V: virulent-virulent.

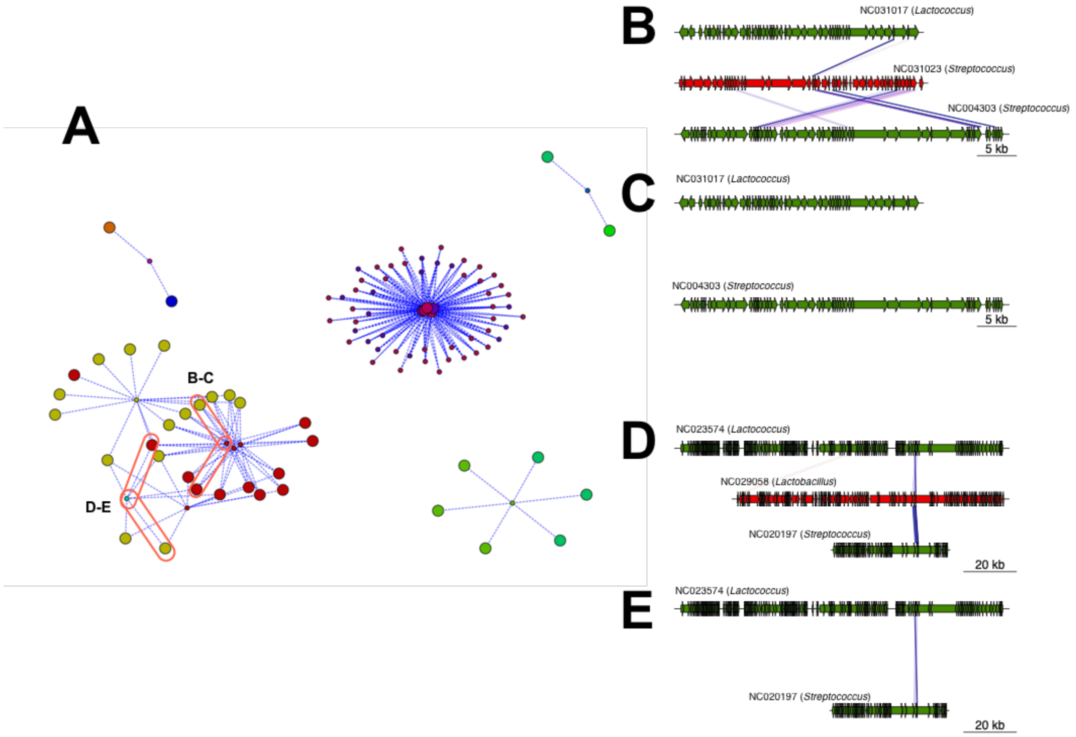


**Fig. S19. Genetic exchanges between temperate phages with different host genera through virulent phages, with the phages' lifestyle inferred with BACPHLIP.** A) Simplified network restricted to pairs of temperate-virulent phages where virulent phage shared genes with at least two temperate phages infecting distinct bacterial genera (i.e., edges shown are only those that link temperate phages with different hosts' genuses through a virulent phage). Recombination data originates from Fig S4. Each node color represents a different bacterial host genus, and the node sizes identify either temperate (large nodes) or virulent (small nodes) phages. Examples from panels B-C and D-E are highlighted in ellipses. **B)** Example of a virulent phage (red) with genes shared with two temperate phages (green) infecting distinct bacterial genera. **C)** Homology (or lack of) between the two temperate phages shown in panel B. D) Analogous to B, but representing exchanges with the virulent phage that involve the exact same protein in both temperate phages**.** **E)** Homology between the two temperate phages shown in panel C. **B-E)** Colors in the blocks linking the phages indicate the level of sequence similarity between homologs.

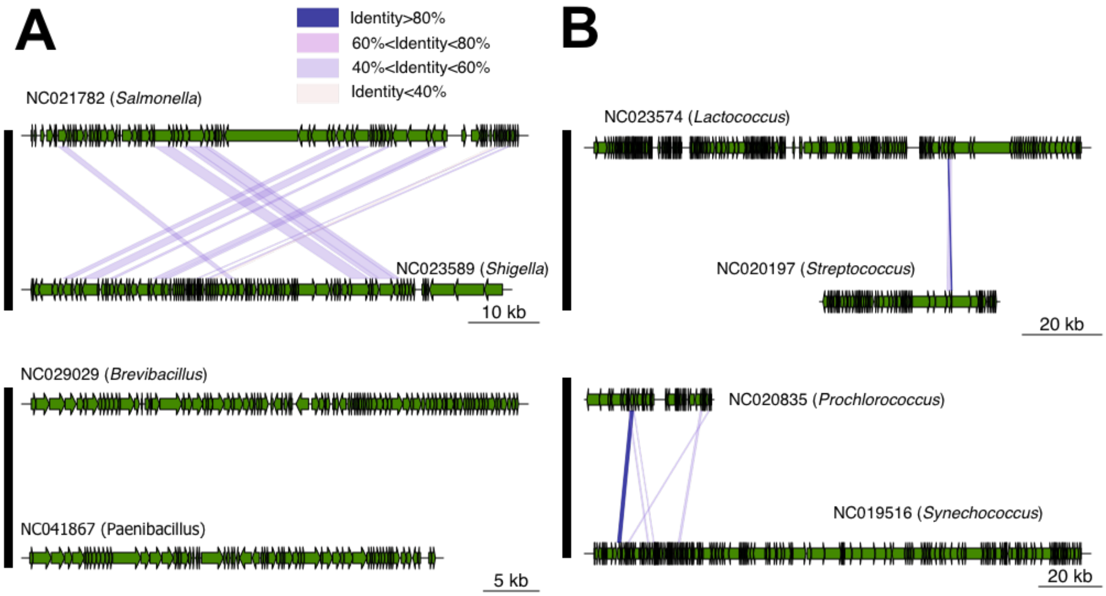


**Fig. S20. Genomic maps of the relationships of homology between temperate phages with different host genera, that are connected through genetic transfers with virulent phages.** The genomes of temperate phages correspond to the ones shown in Figure 5B-C (panels **A** and **B**, respectively) in the main text. Colors in the blocks linking the phages indicate sequence similarity between homologs.

**Table S1. Summary of the main results across multiple methods of classifying the lifestyle of phages.**

| | PHASTER (All phages) | PHASTER (Only phages with confidently assigned lifestyle) | BACPHLIP |
|---|---|---|---|
| ***Lower wGRR between phages with different lifestyles (compared to intra-lifestyle phage pairs)*** | Yes (Fig 1A) | Yes (Fig S1A) | Yes (Fig S1C) |
| ***wGRR clusters more heterogeneous for phage lifestlye (compared phage family or host phyla)*** | Yes (Fig 1B) | Yes (Fig S1B) | Yes (Fig S1D) |
| ***Identification of pairs of recently mosaic dissimilar phages, within and between lifestyles*** | Yes (Fig 2, Fig 3, Fig S8A-B) | Yes, (Fig S5A-B, Fig S8C-D) | Yes (Fig S4, S5 C-D, S9) * & ** |
| ***Diversity of phage-like functions transferred between temperate and virulent phages*** | Yes (Fig 4A and File S2) | Yes (Table S3) | Yes (Fig S12A and File S2) |
| ***Existence of genes with bacteria-like functions transferred between temperate and virulent phages*** | Yes (Fig 4B and File S2) | Yes (Table S4) | Yes (Fig S12B and File S2) *** |
| ***Enrichment of phage-like recombinases in recently mosaic phages*** | Yes (Table S5) | Yes (Table S5) | Yes (Table S5) |
| ***Enrichment of phage-like recombinases in the hosts of recently mosaic phages*** | Yes (Table S7) | Yes (Table S7) | Yes (Table S7) |
| ***Enrichment of NHEJ in the hosts of recently mosaic phages*** | Yes (Fig S16) | Yes (Fig S16) | Yes **** |
| ***Broader host range of recently mosaic virulent phages (compared to recently mosaic temperate phages)*** | Yes (Fig 6, S18) | Yes (Fig S17) | Yes (Fig S17) |
| ***Identification of examples of virulent phages as putative shuttlers of genes between temperate phages with different bacterial host genus*** | Yes (Fig 7, S20) | NA | Yes (Fig S19) ***** |

* **Due to a lack of statistical power, the linear model based on the Temperate-Virulent dataset could not be used to identify significant outliers (i.e., recombinant pairs). Instead, we used the virulent-virulent dataset, which should also be more heavily weighted by ancestry than by gene transfer events. Indeed, we find that the areas of genetic transfers identified with this alternative approach are quite similar to those obtained in the main analysis (using the linear model based on the temperate-virulent phage pairs). All subsequent analysis using the BACPHLIP classification were performed with the dataset of recombinant pairs identified with the virulent-virulent linear model.**

** **Fraction of recently mosaic virulent phages=45%**
**Fraction of recently mosaic temperate phages=77%**
**Fraction of virulent phages with gene transfers with temperate phages=17%**
**Fraction of temperate phages with gene transfers with virulent phages=32%**

*** **Number of genes with bacterial functions is reduced, but still includes genes related with photosynthesis**

******** **Mean NHEJ in hosts of recently mosaic phages=0.406**
**Mean NHEJ in hosts of the other phages=0.359**
**2-sample KS test pval=0.005**

********* **Some of the examples shown in the main text are not verified with the analysis based on BACPHLIP, due to disparaging classifications of individual phages (those examples are still classed as recently mosaic)**

**Table S2. Stepwise logistic regression of the presence/absence of recent genetic transfers between a pair of phages (dependent binary variable) and the lifestyle and the genetic distance (wGRR) of the pair (as independent variables).** The final $R^2$ corresponds to that observed at the final step (#3). The table indicates the BIC and AIC criteria, the former was used as selection criteria. The first analysis concerns pairs where wGRR≥1% whereas for the second wGRR ≥5%. In both cases, the full model (introduction of all variables) was highly significant (p<0.0001), as were the Effect Likelihood Ratio Tests for both variables (always p<0.0001).

| Step | Variable | RSquare | AIC | BIC |
|---|---|---|---|---|
| **wGRR>0.01** | | | | |
| 1 | TT/VT/VV{VV&VT-TT} | 0,0552 | 80706,81 | 80726,07 |
| 2 | TT/VT/VV{VV-VT} | 0,0562 | 80624,12 | 80653,01 |
| 3 | wGRR | 0,0569 | 80566,13 | 80604,65 |
| **wGRR>0.05** | | | | |
| 1 | TT/VT/VV{VT&VV-TT} | 0,1018 | 41769,82 | 41787,23 |
| 2 | wGRR | 0,2212 | 36219,20 | 36245,31 |
| 3 | TT/VT/VV{VT-VV} | 0,2213 | 36216,35 | 36251,17 (NS) |

**Table S3. Bacteriophage functions associated with genetic transfers between temperate and virulent phages, considering only the recently transferred proteins between phages with a confidently assigned lifestyle.** pVOG profiles matching the phage protein dataset (first column). Number of recently transferred proteins (between temperate and virulent phages, second column). Enrichment (+) or depletion (-) of the functional category in recently transferred proteins relative to all phage proteins (Fisher-exact test, significant differences for each functional category shown as asterisks (*) and adjusted for multiple comparisons using the Benjamini-Hochberg method).

| pVOG Category | Matched profiles | Matched recently transferred proteins in temperate-virulent phage pairs | Adjusted significance |
|---|---|---|---|
| ***Unknown*** | 51 | 179 | **** (+) |
| ***Structure*** | 6 | 8 | **** (-) |
| ***DNA metabolism, Regulation & Recombination*** | 11 | 23 | ns |
| ***Packaging, Injection & Assembly*** | 4 | 5 | ** (-) |
| ***Others*** | 1 | 1 | n.s. |
| ***Lysis*** | 2 | 2 | n.s. |
| ***Structure & Lysis*** | 2 | 3 | n.s. |
| ***Other combinations*** | 1 | 1 | n.s. |

**Table S4. Bacterial functions associated with genetic exchanges between temperate and virulent phages, considering only the recently transferred proteins between phages with a confidently assigned lifestyle.** Columns contain the number of bactNog profiles matching the phage protein dataset and number of recently transferred proteins matched by the profiles.

| bactNOG Category | Matched profiles | Matched recently transferred proteins in temperate-virulent phage pairs |
|---|---|---|
| ***Function unknown*** | 23 | 33 |
| ***Replication, recombination and repair*** | 5 | 5 |
| ***Transcription*** | 1 | 5 |
| ***Energy production and conversion*** | 2 | 47 |
| ***Carbohydrate transport and metabolism*** | 0 | 0 |
| ***Inorganic ion transport and metabolism*** | 0 | 0 |
| ***Cell wall/membrane/envelope biogenesis*** | 1 | 1 |
| ***Nucleotide transport and metabolism*** | 1 | 2 |
| ***Transcription + Signal transduction mechanisms*** | 1 | 1 |

**Table S5. Presence of recombinases in recently mosaic and the other phages.** Numbers and proportion of phages with (right column) or without (left column) phage-like recombinases in their genomes. For PHACTS (All), distributions are significantly different both for temperate phages (Fisher exact test, pval<1e-11) and for virulent phages (pval<1e-18). For PHACTS (Confident only), distributions are significantly different both for temperate phages (Fisher exact test, pval<1e-11) and for virulent phages (pval<1e-12). For BACPHLIP, distributions are significantly different both for temperate phages (Fisher exact test, pval<1e-08) and for virulent phages (pval<1e-14).

| | | Recombinases | |
|---|---|---|---|
| Lifestyle classification | Phage class | Absent | Present |
| PHACTS (All) | Temperate Recently Mosaic (445) | 330 (74%) | 115 (26%) |
| | Temperate Other (852) | 493 (58%) | 359 (42%) |
| | Virulent Recently Mosaic (562) | 400 (71%) | 162 (29%) |
| | Virulent Other (528) | 257 (49%) | 271 (51%) |
| PHACTS (Confident only) | Temperate Recently Mosaic (180) | 140 (78%) | 40 (22%) |
| | Temperate Other (587) | 306 (52%) | 281 (48%) |
| | Virulent Recently Mosaic (244) | 162 (66%) | 82 (34%) |
| | Virulent Other (298) | 118 (40%) | 180 (60%) |
| BACPHLIP | Temperate Recently Mosaic (206) | 162 (79%) | 44 (21%) |
| | Temperate Other (707) | 399 (56%) | 308 (44%) |
| | Virulent Recently Mosaic (815) | 580 (71%) | 235 (29%) |
| | Virulent Other (656) | 336 (51%) | 320 (49%) |

**Table S6. Presence of recombinases in Temperate and Virulent HGCF and LGCF phages.**

| | **No recombinases** | **Recombinases** |
|---|---|---|
| Temperate HGCF (249) | 94 (38%) | 155 (62%) |
| Temperate LGCF (227) | 166 (73%) | 61 (27%) |
| Virulent HGCF (34) | 13 (38%) | 21 (62%) |
| Virulent LGCF (510) | 257 (50%) | 253 (50%) |

**Table S7. Average number of phage-like recombinases in the phages' bacterial host species.** Values in bold indicate significantly different distributions (*** p<0.001, * p<0.05, 2 sample Kolmogorov Smirnof test).

| | | Mean number of recombinases | | |
|---|---|---|---|---|
| Lifestyle classification | Host of | Total | Outside prophages | In prophages |
| PHACTS (All) & BACPHLIP | Recently Mosaic Phages | **0.75 (***)** | 0.05 | **0.7 (***)** |
| | Other Phages | 0.58 | **0.054 (*)** | 0.53 |
| PHACTS (Only confidently assigned phage hosts) | Recently Mosaic Phages | **0.75 (***)** | 0.05 | **0.7 (***)** |
| | Other Phages | 0.58 | **0.054 (*)** | 0.53 |
| PHACTS (Confident lifestyle only) | Recently Mosaic Phages | **0.86 (***)** | **0.06 (*)** | **0.8 (***)** |
| | Other Phages | 0.63 | 0.05 | 0.56 |
| PHACTS (Confident lifestyle and with confidently assigned phage host) | Recently Mosaic Phages | **0.94 (***)** | **0.07 (*)** | **0.88 (***)** |
| | Other Phages | 0.71 | 0.06 | 0.65 |

**File S1 (separate file).** List and complete information for all the phage genomes retrieved from RefSeq and used in this study.

**File S2 (separate file).** Detailed listing and description of the phage-like (pVOG) or bacteria-like (bactNOG) profiles matched with the genes transferred within and between phage lifestyles (PHACTS and BACPHLIP).

**File S3 (separate file).** Detailed analysis of recombinases, NHEJ and IS per phage lifestyle (PHACTS and BACPHLIP).

**File S4 (separate file).** Detailed listing of patristic distance and genomic distance for the phage pairs analysed in Fig 6 of the main text (PHACTS only).

**File S5 (separate file).** Detailed pVOG functions and their classification into major functional categories.

**File S6 (separate file).** Hidden Markov Models for the sak4 and gp2.5 recombinases.
